# Long-read transcriptomics of a diverse human cohort reveals widespread ancestry bias in gene annotation

**DOI:** 10.1101/2025.03.14.643250

**Authors:** Pau Clavell-Revelles, Fairlie Reese, Sílvia Carbonell-Sala, Fabien Degalez, Winona Oliveros, Carme Arnan, Roderic Guigó, Marta Melé

## Abstract

Accurate gene annotations are fundamental for interpreting genetic variation, cellular function, and disease mechanisms. However, current human gene annotations are largely derived from transcriptomic data of individuals with European ancestry, introducing potential biases that remain uncharacterized. Here, we generate over 800 million full-length reads with long-read RNA-seq in 43 lymphoblastoid cell line samples from eight genetically-diverse human populations and build a cross-ancestry gene annotation. We show that transcripts from non-European samples are underrepresented in reference gene annotations, leading to systematic biases in allele-specific transcript usage analyses. Furthermore, we show that personal genome assemblies enhance transcript discovery compared to the generic GRCh38 reference assembly, even though genomic regions unique to each individual are heavily depleted of genes. These findings underscore the urgent need for a more inclusive gene annotation framework that accurately represents global transcriptome diversity.

## Main

The accurate annotation of the human genome is crucial for understanding the molecular mechanisms underlying cellular function in health and disease. Gene annotations capture the multiple transcript isoforms produced from each gene by alternative splicing, transcription initiation and polyadenylation. These can alter the transcript’s fate or change its functional product^1–5^. Gene annotations are necessary to understand genome function as they provide essential context for the interpretation of omics measurements (RNA-seq, ChIP-seq, ATAC-seq, etc.), and are critical to the mechanistic understanding of genetic variation. To enable unbiased investigation of health and disease across human populations, these annotations should be representative of the transcriptomic diversity in the human species. Yet, the vast majority of transcriptome data available, from which current reference gene annotations largely derive, comes from biological samples from individuals of European descent^6–9^.

Recently, long-read RNA sequencing (LR-RNA-seq) technologies have transformed the transcriptomics field by enabling unambiguous elucidation of complete transcript structure, as they have the capacity to span the full length of RNA molecules^10,11^. LR-RNA-seq enormously facilitates the comprehensive gene annotation of genomes. The first large-scale efforts to employ full-length transcriptome profiling across human tissues and cell lines have already led to the discovery of thousands of novel transcripts and provided new insights on transcriptome composition^5,12^. Furthermore, employing LR-RNA-seq for the purpose of gene annotation curation has resulted in a massive expansion on the current number of human annotated genes and transcripts^13^. However, all these major LR-RNA-seq studies and annotation efforts have again almost exclusively profiled biological samples obtained from European individuals^5,12^. Therefore, many transcripts exclusive to or more abundant in non-European populations are likely to be underrepresented in human reference annotations (*i.e.,* GENCODE and RefSeq).

Furthermore, current human gene annotations are based on GRCh38, which is a haploid linear reference that inherently lacks genetic variability. This implies that small variants and large structural variation or unique genomic regions, which may harbor novel genes^14^ and/or gene duplication events^15^, are missing from the current assembly. Therefore, reference gene annotations might also remain incomplete due to the choice of the genome assembly. The pressing need for an improved representation of human genetic diversity has resulted in the development of the human pangenome assembly, a graph built using multiple genomes from a cohort of population-diverse samples^16^. The pangenome significantly improves upon the current linear haploid mosaic GRCh38 reference^17^ by representing human genetic variability including single nucleotide polymorphisms (SNPs), short indels, and structural variants^16^. It is expected to advance variant discovery, population genetics studies, disease research, and personalized medicine. However, to what extent transcript discovery and annotation will improve when using personal genome assemblies^18^ is unknown.

In this study, we present the first LR-RNA-seq dataset from a population diverse cohort and provide a comprehensive characterization of the full-length transcriptomes of eight genetically-distinct human populations from lymphoblastoid cell lines (LCL) for a total of 43 samples. Six of these samples have personal genomes available and an additional 30 were genotyped as part of the 1000G project^19,20^. We build a population-diverse gene annotation including more than 30k transcripts never identified before. We show that current reference gene annotations are worse at representing transcripts from non-European populations in comparison to European populations. We identify a subset of 2.2k transcripts that are discovered specifically in a single population and show that they are enriched in novel transcripts only in non-European samples. We demonstrate that this widespread ancestry bias negatively affects our ability to link genetic variants to alternative transcript usage, especially in non-European populations. Then, we quantify the impact of SNPs, which can be interpreted by computational tools as errors, on transcript discovery and show that hundreds of transcripts per sample are potentially missed without accounting for this genetic variation. Similarly, we show that using personal genome assemblies, which contain all facets of genetic variation, enables the discovery of transcripts impossible to detect with GRCh38. Finally, we found that genomic regions not found in linear reference genomes (GRCh38 and T2T) show little transcriptional activity, despite accounting for about 150 Mb per genome (5% of GRCh38 total length). Our study offers new insights into how ancestry-specific biases have impacted current gene annotations and underscores the importance of moving toward inclusive reference annotations for advancing biomedical research equally across diverse human populations.

## Results

### A population-diverse long-read RNA-seq dataset enables the discovery of many high-confidence novel transcripts

We generated long-read RNA-seq (LR-RNA-seq) for 43 genetically-diverse lymphoblastoid cell lines (LCLs) using Oxford Nanopore Technologies (ONT) (Fig. 1a). We used CapTrap^21^ to enrich our samples for complete molecules (Fig. 1b; Methods). Altogether, we profiled samples from eight human populations across four continents: Africa (3), Asia (2), America (1), and Europe (2) (Methods). Our design is sex-balanced and contains between 4 and 6 samples per population, including 6 samples overlapping the human pangenome^16^ and 30 from the 1000G project^19^ (Supplementary Fig. S1; Supplementary Table S1). Overall, we generated 800M reads; one of the most deeply-sequenced LR-RNA-seq datasets to date. After preprocessing and filtering, we retained 8-24 million high-quality reads per sample (Supplementary Fig. S2; Methods).

**Fig. 1:**
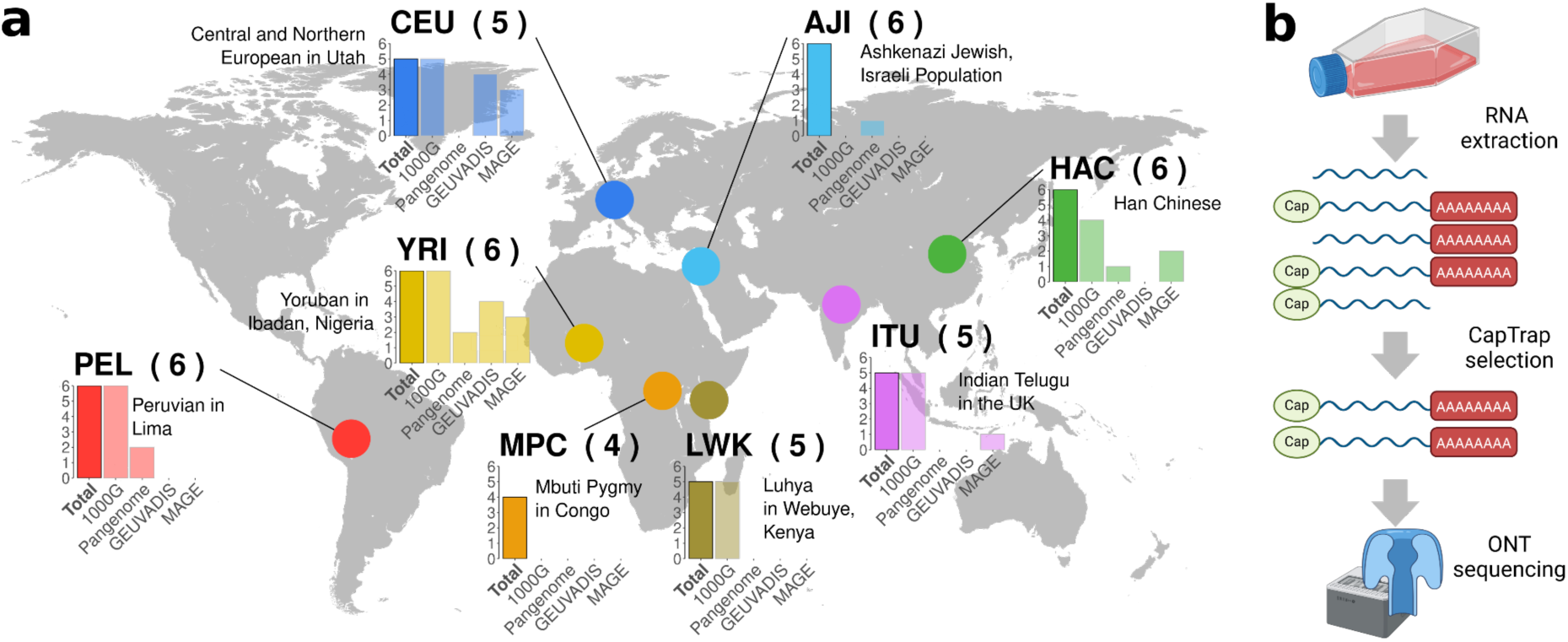
Long-read RNA-sequencing of a diverse population cohort. **a**, Overview of the genetic ancestry of the samples used in this study, the corresponding population name, and whether these have been used in other major genetic diversity projects: the 1000G Project^19,20^, the Human Pangenome Reference Consortium (HPRC)^16^, the GEUVADIS^22^ and the MAGE^23^. **b**, Overview of CapTrap and Oxford Nanopore LR-RNA-seq experimental protocol.

To call transcript models from the filtered, high-quality reads we employed four previously benchmarked^11,24^ tools: FLAIR^25^, IsoQuant^26^ and ESPRESSO^24^, which are annotation-guided (GENCODE v47); and LyRic^27,28^, which is annotation-agnostic. We merged transcript models across samples and tools based on intron chain identity (*i.e.*, transcript models differing only in the ends were considered identical; Methods), such that each transcript corresponds to a unique intron chain. As a consequence, all monoexonic transcripts were removed. The number of distinct transcripts per sample varies from ∼50k-80k and is highly correlated with its sequencing depth (Supplementary Fig. S3; Methods). We obtained a set of 380,519 unique transcript models and discarded 59% based on stringent filtering criteria that varied depending on their gene biotype and SQANTI structural category with respect to GENCODE v47 (Fig. 2b; Supplementary Fig. S4-6; Supplementary Table S2-3; Methods). Full-splice match (FSM) transcripts completely match an annotated intron chain while incomplete-splice match (ISM) transcripts represent an annotated intron chain that is truncated either at the 5’ or 3’ end with respect to an annotated transcript. Novel intron chains are divided into novel in catalog (NIC) and novel not in catalog (NNC) depending on whether they use only annotated splice sites or add novel splice sites respectively. From the discarded transcripts, 76% did not meet multiple criteria. This emphasizes the complementarity of the filters applied and underscores the reliability of the resulting transcript set (Supplementary Fig. S7). The 155,875 transcripts passing these filters compose our final cross-ancestry transcript set, which we call the PODER (POpulation Diversity-Enhanced long-Read) annotation (Supplementary Table S4-5).

**Fig. 2:**
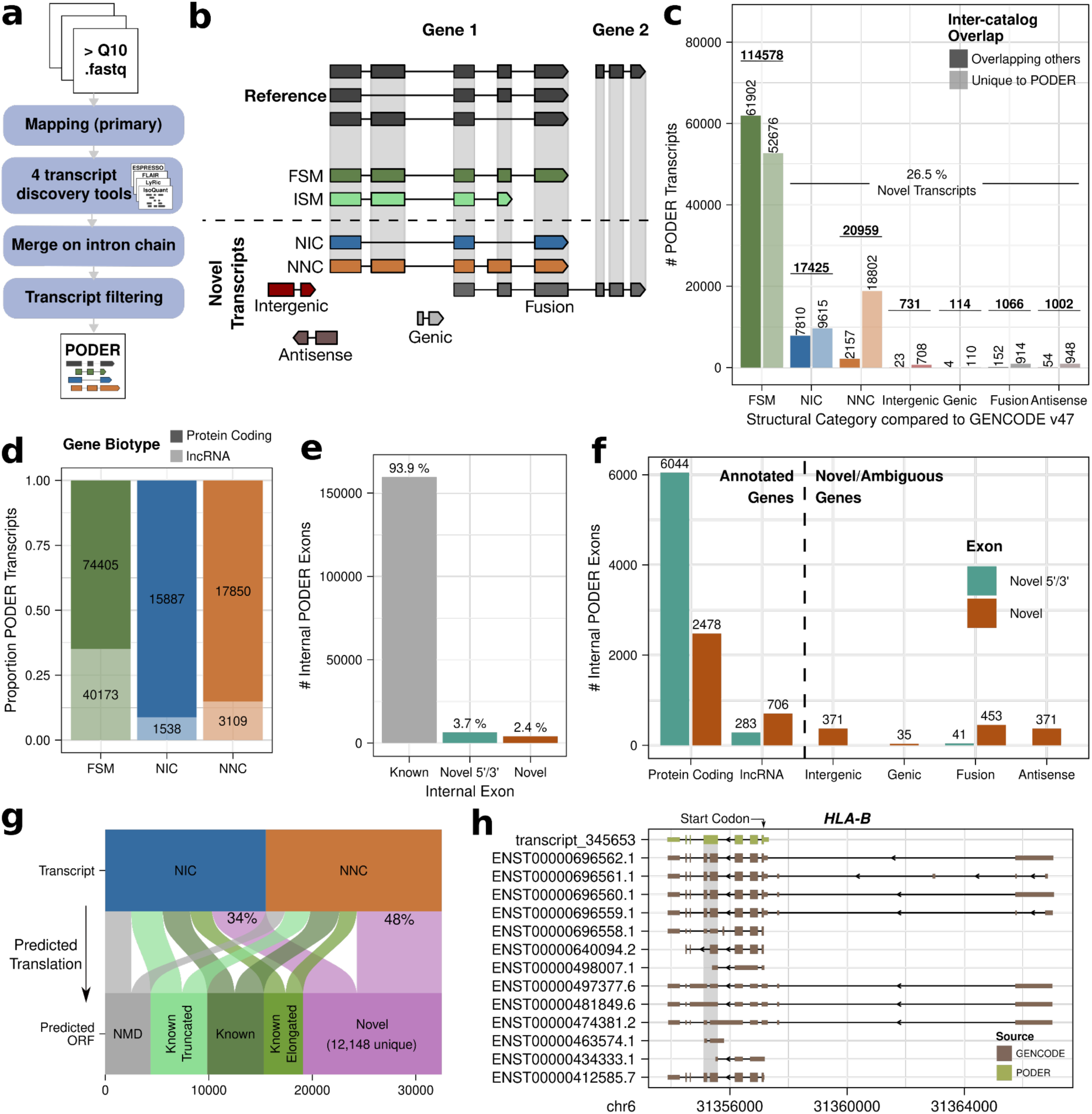
Our population-diverse long-read based gene annotation significantly expands reference gene annotations. **a**, Brief overview of data processing and filtering steps to generate the PODER annotation. Transcript discovery tools include FLAIR, IsoQuant, ESPRESSO and Lyric. **b,** Structural categories of discovered transcripts based on comparison of intron chains (sequence of splice junctions) to the transcripts from a reference gene annotation using the SQANTI classification. FSM: full-splice match, ISM: incomplete-splice match, NIC: novel in catalog, NNC: novel not in catalog (Methods). **c**, Number of PODER transcripts per SQANTI structural category in PODER, stratified depending on if they are also found in another transcript catalog or not (GTEx long-reads v9, ENCODE4, CHESS3). FSMs include 10,255 GENCODE transcripts for which our pipeline called a matching subchain (ISM) that we replaced with its corresponding full annotated structure (Methods). Bold, underlined numbers show the total number of transcripts within each structural category. **d**, Proportion of PODER transcripts gene biotypes by SQANTI structural category. **e**, Number and percentage of internal exons by annotation status in GENCODE. Novel 5’/3’ refers to 5’ or 3’ extensions or truncations of previously annotated exons. **f**, Number of novel internal exons by gene biotype or SQANTI structural category if belonging to novel/ambiguous genes. Ambiguous genes are those that contain fusion transcripts spanning two different gene bodies in the same strand. **g**, Correspondence between structural categories of novel transcripts from protein coding genes and their predicted ORF status (Methods). NMD (nonsense mediated decay) means ORF is predicted to be non-productive, “known” means annotated in GENCODE, “known truncated” is a subsequence of an annotated protein sequence, “known elongated” represents extensions of amino acids of annotated protein sequences, and “novel” includes at least one amino acid difference from annotated sequences that is neither a truncation or elongation (Methods). Different transcripts can have the same predicted ORF, therefore predicted ORF bins can include redundant sequences (*i.e.,* there are 12,148 unique novel amino acid sequences predicted from 13,401 transcripts). **h**, Example of a novel transcript with a novel predicted amino acid sequence. GENCODE *HLA-B* transcripts (brown) and a novel (NIC) PODER transcript (green). Novel PODER transcript exhibits a frame-preserving intron retention event (shaded in gray). CDSs (predicted for novel transcripts and annotated for GENCODE transcripts) are indicated by thicker regions within transcript structure.

Among PODER transcripts, 73.5% are already annotated in GENCODE, and thus 41,293 PODER transcripts are novel (Fig. 2c). Most novel transcripts belong to protein-coding genes, which is expected given that such genes are more highly-expressed compared to lncRNAs^29,30^ (Fig. 2d; Supplementary Fig. S8-9; Supplementary Results; Methods). We compared PODER transcripts with those discovered by recent large-scale LR-RNA-seq transcriptome studies from GTEx^12^ and ENCODE4^5^; as well as with those derived from short-read RNA-seq data from CHESS3^31^. These catalogs show similar rates of overlap between each other for novel transcripts (20-35%; Supplementary Fig. S10). Despite sequencing only a single cell line, we discover the second highest number of FSM transcripts compared to the rest of catalogs, which are all more complex in terms of sample composition. This demonstrates our high sensitivity for annotated transcripts. Furthermore, we discover 31,097 novel transcripts (75.3%) that have not already been described by the other catalogs (Fig. 2c; Supplementary Fig. S10a; Supplementary Table S6; Supplementary Results).

Novel PODER transcripts include 10,785 novel internal (*i.e.,* not terminal) exons which either overlap already-annotated exons and use novel splice sites (Novel 5’/3’; 6,327) or are entirely novel (Novel; 3,184; Fig. 2e-f; Methods). The latter are more common in transcripts from lncRNAs and novel genes whereas the former are more abundant in protein-coding genes (Fig. 2f). Novel exons from PODER contain SNPs with greater allelic divergence between CEU and non-European populations (YRI, LWK, HAC, PEL, ITU) than known exons, demonstrating that differences in allele frequencies between European and non-European populations might influence exon annotation status (Supplementary Fig. S11; Methods). We evaluated the coding potential of novel transcripts from protein-coding genes and found that 41% of them are predicted to encode for a novel amino acid sequence which does not represent a simple extension or truncation of an existing sequence (Fig. 2g-h; Methods). Finally, we clustered novel intergenic transcripts into 476 unique novel genes, representing parts of the genome with novel transcription undergoing splicing (Supplementary Fig. S12a; Methods). Novel genes have fewer transcripts per locus (1.53 on average), the shortest transcript lengths, and the lowest number of exons per transcript (Supplementary Fig. S12). Adding the novel PODER transcripts to the GENCODE annotation (to create the Enhanced GENCODE; Supplementary Table S7) increases the number of reads contributing to gene expression quantification (152,787, 0.67%; median number and percent increase, respectively) using the 731 MAGE^23^ LCL short-read RNA-seq samples (Supplementary Table S8-10; Methods). In addition, predicted variant effects with VEP^32^ on Enhanced GENCODE have similar proportions of impact severity to those predicted on GENCODE alone (Supplementary Fig. S13; Supplementary Results; Methods).

Overall, we produced and curated the first human gene annotation using deeply-sequenced population-diverse LR-RNA-seq samples, which enabled us to discover over 30k transcripts never identified before.

### Current gene annotations are less representative of transcriptomes from non-European descent individuals

To assess the possibility of a European ancestry bias in the annotation, we first classified our samples into European (CEU, AJI) and non-European (YRI, LWK, MPC, HAC, PEL, ITU), although we acknowledge the limitations of such broad and discrete categorizations (Supplementary Table S1; Methods). When comparing the number of transcripts discovered in each sample, we found a significantly higher number of novel PODER transcripts discovered in non-European samples (Fig. 3a; Supplementary Fig. S14, Supplementary Results). This bias is not the consequence of higher genetic diversity in African populations^19^, as we did not observe differences between African and out-of-Africa (OOA) populations in the number of novel transcripts discovered (Supplementary Fig. S14). Thus, non-European transcripts are less represented in reference gene annotations.

**Fig. 3:**
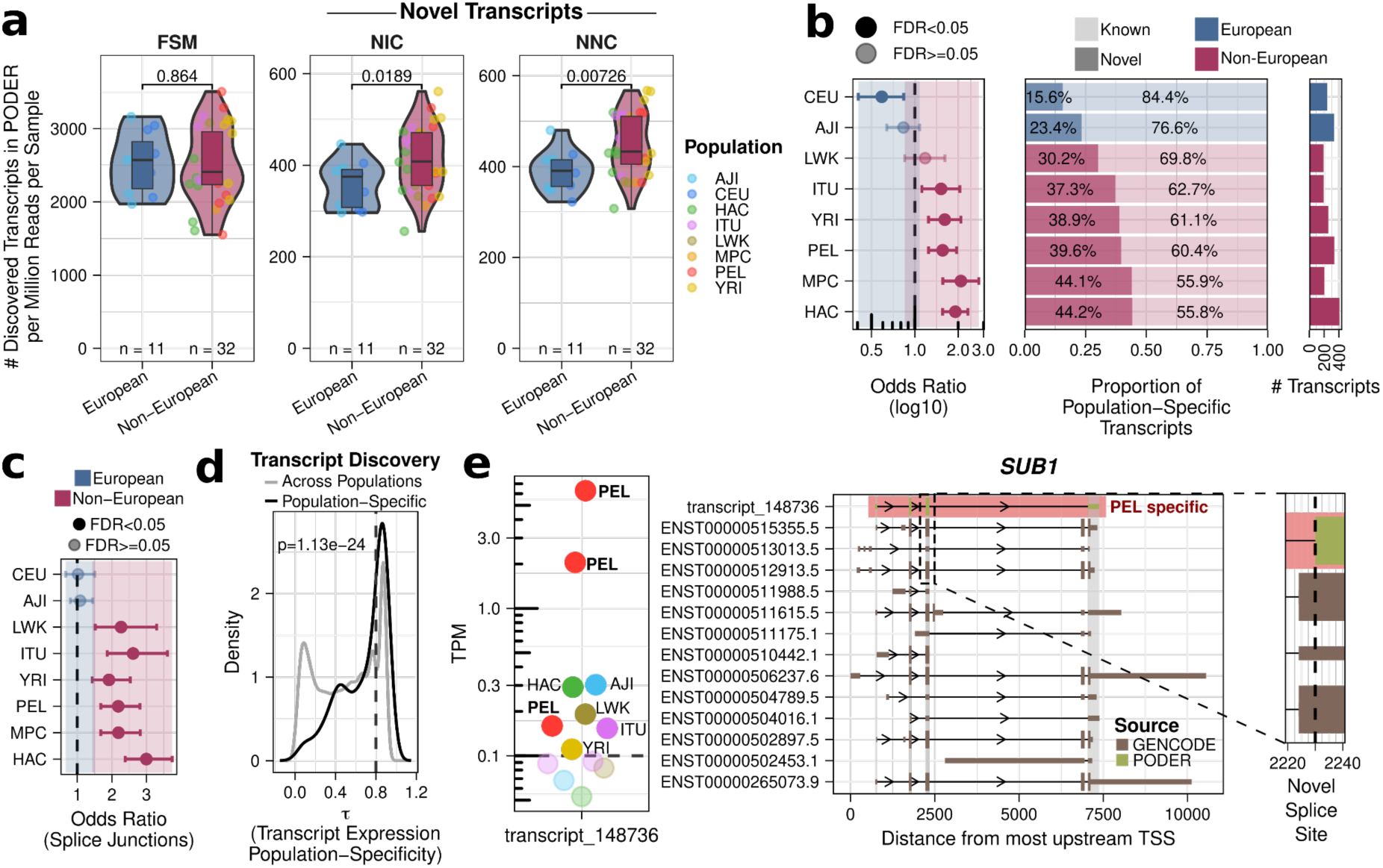
Population diverse full-length transcriptomes reveals European bias in reference gene annotation. **a,** Number of PODER transcripts discovered per sample normalized by sequencing depth between European vs. non-European populations by structural category (t-test, FDR adjusted by Benjamini–Hochberg). **b**, Enrichment in novel transcripts of population specific transcripts per population (left). Proportion of population-specific transcripts by annotation status (middle). Number of population-specific transcripts per population (right). **c**, Enrichment in novel splice junctions per population. **d,** Tau quantification-based population-specificity scores of transcripts, split by population-specificity on discovery. Line at 0.8 indicates threshold of high expression-based population-specificity. **e**, Example of population-specific discovered transcript (NNC) from gene *SUB1* in Peruvians (PEL) containing a novel splice site in the third exon (5’ exon shortening) producing a predicted in-frame modification of the CDS. Left: transcript expression per sample in TPM. Middle: annotated (brown) and population-specific novel (green) transcript models. CDSs (predicted for novel transcripts and annotated for GENCODE transcripts) are indicated by thicker regions within transcript structure. Right: magnification of novel splice site.

We wondered how the different populations in our dataset enable the differential discovery of certain transcripts. Therefore, we classify transcripts and splice junctions as population-specific if they were discovered in at least two samples exclusively in a single population (Methods). These are transcripts that would have been missed during transcript discovery if we had not included their population in this study, although they might also exist in other populations at levels that impede their detection during transcript discovery. We identified 2,267 population-specific transcripts (population median 260; 1.46% of transcripts from PODER). Population-specific transcripts are enriched in lncRNAs, are significantly more lowly-expressed and shorter than population-shared transcripts (Supplementary Fig. S15).

European population-specific transcripts are significantly enriched in already-annotated transcripts compared to non-European population-specific transcripts, which are enriched in novel ones (Fig. 3b; Supplementary Fig. S16a). This trend is reproducible when performing uniform read depth downsampling, when using RefSeq instead of GENCODE as the reference annotation, and is agnostic of our filtering strategies (Supplementary Fig. S16; Supplementary Results; Methods). Accordingly, population-specific splice junctions are also enriched in novel splice junctions only in non-European populations (Fig. 3c; Supplementary Fig. S15a). Overall, there is more novelty in population-specific transcripts from non-Europeans than in population-shared transcripts, demonstrating that the effect of the ancestry bias in gene annotations is exacerbated when considering transcripts that are discovered specifically in one certain population.

Our definition of population specificity based on transcript discovery is a technical, discrete categorization inherently limited by the composition of our cohort and partially influenced by the underlying biological phenomenon of expression. Therefore, we used the Tau (τ) specificity metric to assess how population-specific discovery relates to the underlying continuum of transcript expression across ancestries (Supplementary Table S5, S11-12; Methods)^33^. As expected, population-specific discovered transcripts have significantly higher τ specificity values than shared transcripts discovered across populations, even in the aforementioned MAGE dataset when quantified with Enhanced GENCODE (Fig. 3d-e; Supplementary Fig. S17; Supplementary Table S13). Furthermore, we demonstrate that the 8,740 highly population-specific expressed transcripts (τ ≥ 0.8; 4,154 of which are novel) are significantly enriched in population-specific transcripts (Fisher’s test, p-value < 2·10^-16^). Again, this finding replicates when quantifying the short-read population-diverse MAGE RNA-seq dataset; demonstrating its robustness in the context of a higher sample size and more distinct human populations (Supplementary Fig. S17; Supplementary Results; Methods). In conclusion, these results suggest that human reference gene annotations are depleted of non-European transcripts at both the global and population-specific level, which cannot solely be explained by differences in genetic diversity between populations.

### A population diverse-annotation enhances discovery of allele-specific transcript usage in non-European populations

It has been shown that leveraging population diverse cohorts improves the discovery and fine-mapping of genetic effects on transcriptomic traits^14,23,34^. Our LR-RNA-seq dataset is especially suited to identify such genetic effects impacting transcript usage as we can assign full transcripts to specific alleles. Thus, we performed allele-specific expression (ASE) and allele-specific transcript usage (ASTU) analyses on the subset of 30 of our LR-RNA-seq samples with available phased genotype data (CEU, HAC, ITU, LWK, PEL, YRI)^20^. ASE associates genetic variants with differences in gene expression between alleles whereas ASTU identifies allele-specific impacts on transcript isoform usage (Fig. 4a).

**Fig. 4:**
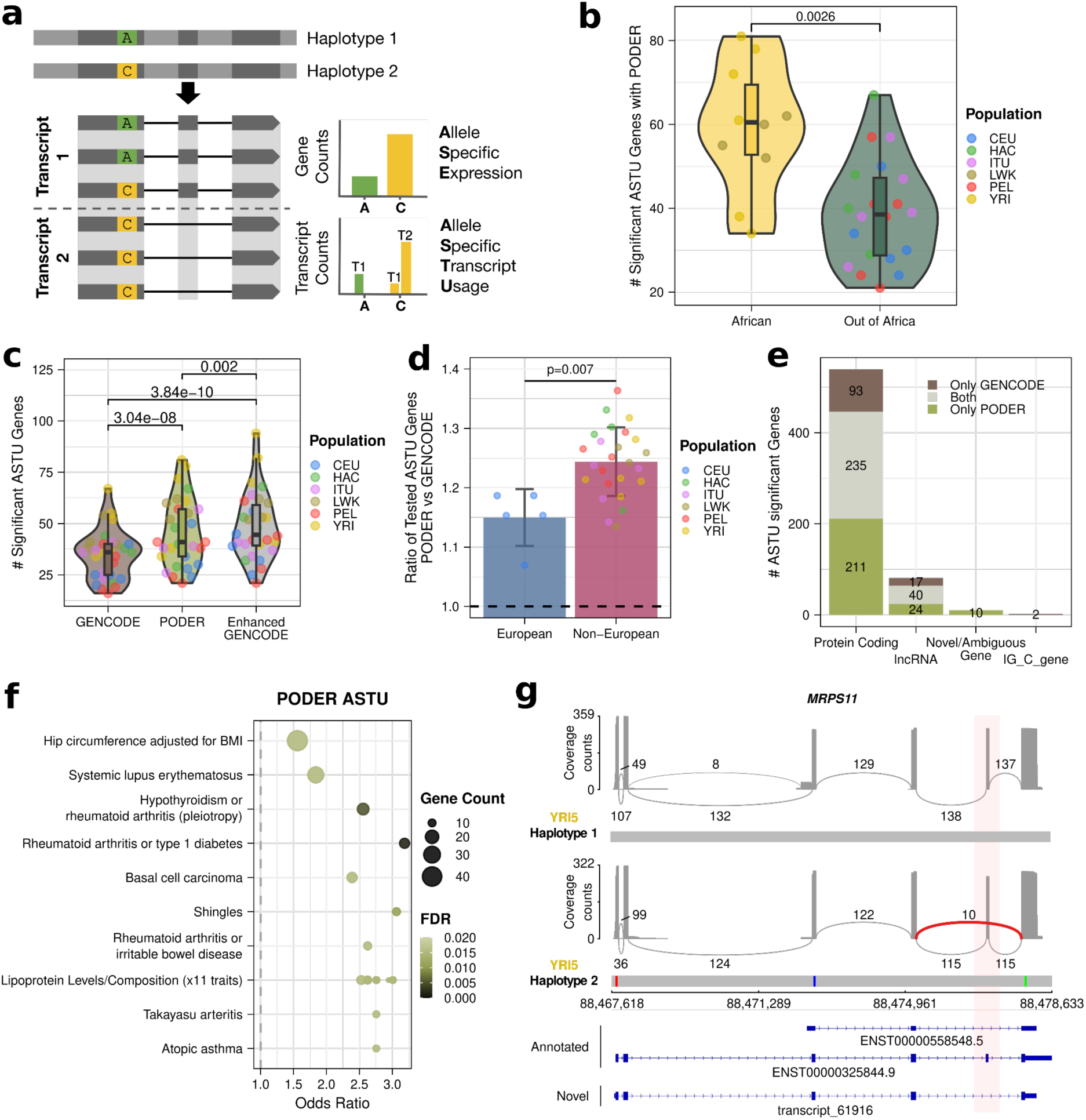
A genetically diverse annotation increases the potential for discovery of allele-specific transcript usage in non-European populations. **a**, Overview of allele-specific expression and transcript usage (ASE and ASTU respectively)^12^. **b**, Number of ASTU significant genes in African and out of Africa (OOA) samples (t-test) with PODER. **c**, Number of ASTU significant genes per sample when using GENCODE, PODER and Enhanced GENCODE (paired t-test, FDR corrected). **d**, Ratio of number of tested genes when using PODER versus GENCODE for each sample, by population (t-test). **e**, Number of ASTU significant genes shared between PODER and GENCODE, by gene biotype. **f**, PODER ASTU significant genes enrichment in GWAS hit associated genes (cutoff FDR = 0.02) **g**, Sashimi plots demonstrating junction use frequency (arcs) and read coverage (y-axis) in the *MRPS11* allele-specific transcript usage gene per haplotype for sample YRI5. Only junctions with ≥ 8 reads are shown. Representative haplotypes (exonic only) shown. There are no GENCODE transcripts that are long and skip this exon, therefore transcript models from GENCODE were chosen to demonstrate this. Novel transcript model from PODER chosen to show full model with the exon skipping event.

To assess the relative benefit of incorporating novel transcripts from our cohort into standard annotations, we performed these analyses thrice using different references: PODER, GENCODE, or Enhanced GENCODE. Using PODER, we found a median of 278 ASE and 50 ASTU significant genes per sample, which are mainly protein-coding (Supplementary Fig. S18-19; Supplementary Table S14-15). 53.5% of ASE and 66.5% of ASTU significant genes are reproducible in at least two samples, demonstrating reproducible genetic control of gene expression and alternative transcript usage (Supplementary Fig. S20). We discover slightly more ASE and ASTU genes per sample than previous efforts^12^, likely thanks to improvements in sequencing quality, our higher sequencing depth, and the higher genetic diversity of our samples. Both ASE and ASTU are significantly enriched in e/sGenes defined in LCLs from the GTEx project^35^ (Fisher’s test, p-value = 2.2·10^-15^ and 1.4·10^-25^, respectively), and show a lower overlap for Africans than OOA populations, likely due to the proportionally lower number of African-descent samples in GTEx^35^ (Wilcoxon’s rank-sum, p-value = 0.00032-0.037; Supplementary Fig. S21). As expected, African individuals have more tested and significant ASE and ASTU genes than individuals from OOA populations, concordant with Africans having higher heterozygosity^36^ (t-test, p-value = 0.0016-0.04; Fig. 4b; Supplementary Fig. S22; Methods).

An annotation derived from the transcriptome of an ancestry-diverse cohort like PODER is more representative of population diversity and could increase the discovery of genetic effects on gene expression and transcript usage. We find that the number of ASE genes discovered with PODER, GENCODE, and Enhanced GENCODE annotations is comparable (Supplementary Fig. S23a). However, both PODER, and even more so Enhanced GENCODE, test more genes for ASTU and accordingly identify more ASTU-significant genes compared to GENCODE alone (Fig. 4c; Supplementary Fig. S23b). The fact that we gain ASTU genes using both PODER and Enhanced GENCODE demonstrates that novel transcripts increase ASTU discovery by completing existing annotations rather than providing tailored ones specific to the dataset. Furthermore, the relative increase in number of ASTU-tested genes in PODER versus GENCODE is significantly higher in all non-European populations than in Europeans (CEU), with a mean increase of 1.15x in Europeans and 1.24x in non-Europeans (Fig. 4d). The increase in testability is directly attributable to the addition of novel, expressed transcripts to genes which previously only had one expressed transcript (Supplementary Results; Supplementary Fig. S24; Methods). This demonstrates that while all ancestries benefit from a more complete annotation, the European bias in gene annotations disproportionately impacts non-Europeans in the discovery of genetic effects on transcript usage.

Genetic variation influencing gene expression or transcript usage might be key to understanding population-level disparities in disease susceptibility^37,38^. To explore this, we investigated whether our identified ASE and ASTU genes were enriched in genes associated with GWAS traits from the GWAS catalog^39^. While ASE genes from both PODER and GENCODE had no significant enrichments, ASTU genes were enriched in several non-redundant GWAS traits (6 with GENCODE and 10 with PODER; Fig. 4f; Supplementary Fig. S25; Fisher’s test, FDR ≤ 0.02; Supplementary Table S16; Methods). Systemic lupus erythematosus^40^, Takayasu’s arteritis^41^, and rheumatoid arthritis^42^ are autoimmune diseases and along with basal cell carcinoma^43^ and atopic asthma^44^, have a different incidence or severity between different human populations^40,45,46^. Additionally, PODER-specific ASTU genes are enriched in GWAS traits related to levels and composition of cholesterol and lipoproteins which also present known differences between ancestries^47,48^. Our findings underscore the importance of incorporating novel transcripts from diverse populations into gene annotations, as this approach enhances the sensitivity of genetic effect discovery and can provide deeper insights and interpretation into population-specific differences in disease susceptibility.

### Personalized genome reference assemblies enhance novel splice junction discovery

LR-RNA-seq transcript discovery tools employ various strategies to filter false positive splice junctions that arise due to common LR-RNA-seq artifacts such as miscalled bases and microindels^49^. These include enforcing novel splice site motifs to be canonical (GT/AG, GC/AG, AT/AC)^24,50,51^ and requiring perfect matches in alignments close to novel splice junctions^24,26^. However, these requirements will impair the discovery of novel splice junctions when the endogenous haplotypes from the sample and the reference genome sequences (GRCh38) do not match due to genetic variation (Fig. 5a)^52^.

**Fig. 5:**
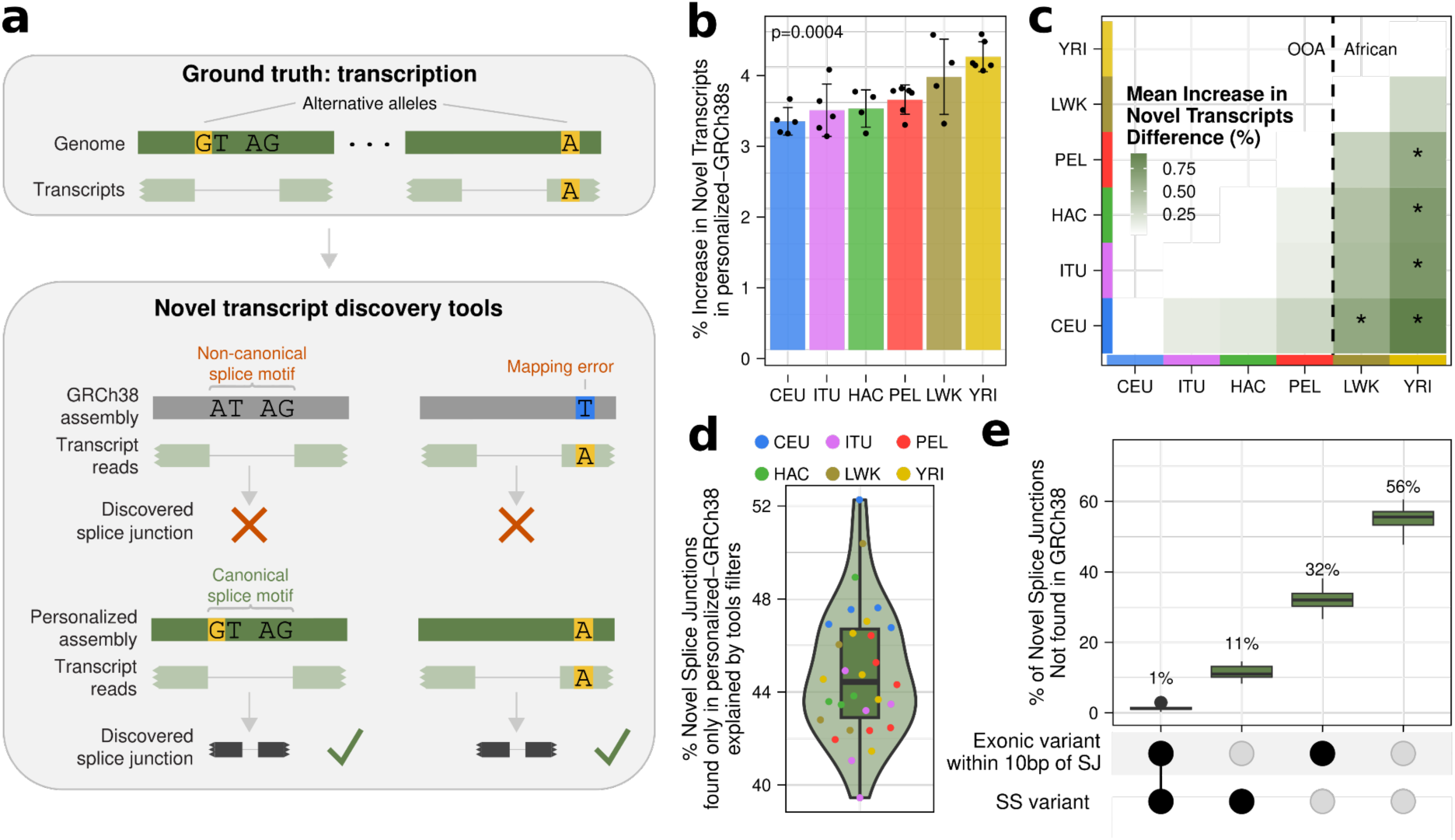
The use of personalized-GRCh38s assemblies uncovers hundreds of novel splice junctions missed with GRCh38. **a,** Common LR-RNA-seq transcript discovery tools restrictions for splice junction discovery. Ground truth transcription from the genome of an individual (top) and filters used in the computational discovery of novel transcripts: 1) presence of a canonical splice site motif in the reference genome assembly and 2) perfect read alignment to the genome in the splice junction vicinity (bottom). **b,** Mean increase (in %) of novel transcripts that are discovered when using the personalized-GRCh38s with respect to the number of novel transcripts discovered using GRCh38, by population. Error bars show standard deviation. Differences tested with an ANOVA. **c,** Post-hoc Tukey test between all populations. Color fill shows mean difference in percentage of transcripts discovered exclusively in personalized-GRCh38s and * means adjusted p-value < 0.05. **d,** Percentage of novel splice junctions identified only in personalized-GRCh38 and not in GRCh38 that can be explained by transcript discovery tool filtering (false negatives). **e,** Upset plot detailing how novel, personalized-GRCh38 unique splice junctions (SJs) are explainable based on presence of genetic variants in splice sites or splice junction-proximal regions.

To quantify this, we ran mapping and transcript discovery in our LR-RNA-seq data using personalized versions of GRCh38 (henceforth personalized-GRCh38s) as the reference genome assembly. For each of our 30 LR-RNA-seq samples with available phased genotype data^20^, we produced two personalized-GRCh38s containing the biallelic SNPs of each phased haplotype (Methods). Using the personalized-GRCh38 haplotypes leads to the detection of a median of 361 (0.83%) more transcripts and 496 more splice junctions (0.55%) per haplotype and sample than using GRCh38 (Supplementary Table S17). Considering only transcripts with novel splicing (everything but FSMs and ISMs), we discover a median of 607 (3.6%) more novel transcripts per sample using the union of the two personalized-GRCh38 assemblies per sample compared to GRCh38 (Fig. 5b). Notably, the increase in novel transcript discovery differs between populations, with African populations exhibiting the largest (ANOVA, p-value=0.0004; Fig. 5b-c). This is concordant with the higher number of alternative alleles in African genomes^19^. For a median of 44.4% (205.5) of novel splice junctions found with personalized-GRCh38s but not discovered with GRCh38, we can directly attribute their non-discovery in GRCh38 to alternative alleles located in splice sites or exonic sequences proximal to splice junctions (within 10bp) (Fig. 5d; Methods). The remaining unexplained discrepancies might be caused by more complex variation not represented in this analysis such as indels. This analysis demonstrates that lack of genetic variation information in reference assemblies intrinsically limits transcript discovery and thus has a larger impact in African descent individuals.

### Personal genome assemblies offer unique advantages for LR-RNA-seq transcript discovery

Genetic variation with respect to GRCh38 consists not only of SNPs but also insertions, deletions, and, most notably, structural variants^19^. Therefore, we investigated the advantages of using full personal genome assemblies containing all such facets of genetic variability rather than GRCh38 for LR-RNA-seq mapping and transcript discovery. We leverage the personal genome assemblies (from now on personal assemblies) available for the six pangenome samples overlapping our LR-RNA-seq dataset^16^. We mapped our LR-RNA-seq data from a given sample to its corresponding personal assembly and to GRCh38 (Methods). On average, 564,640 reads (3.1%) per sample do not map to either genome while 96% of reads per sample map to both (Fig. 6a). Of these, 15% on average map better to the personal assembly than to GRCh38 based on alignment coverage and identity, whereas only 5% map worse (Fig. 6b; Supplementary Fig. S26; Methods). Very few reads exclusively map to personal assemblies (median 1,415; 0.008%) or exclusively to GRCh38 (median 4,847; 0.027%; Fig. 6a). This suggests that personal assemblies enable slightly better mapping qualities than GRCh38 and that few transcriptionally-active genomic regions are exclusive to a single assembly.

**Fig. 6:**
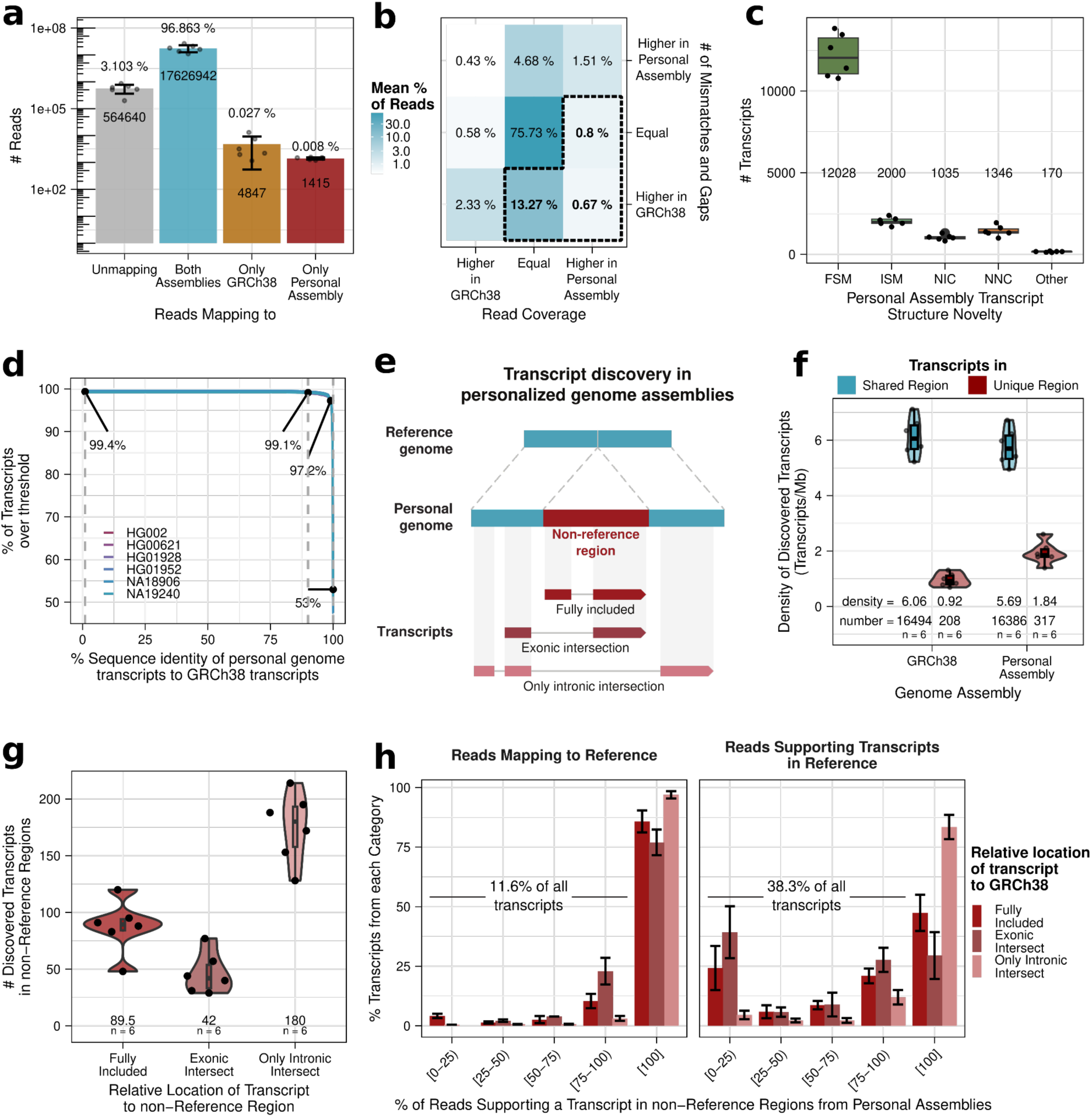
Personal genome assemblies demonstrate benefits for LR-RNA-seq transcript mapping and discovery. **a**, Mean number of reads by the combinations of assemblies where they map to. **b,** Mean percentage of reads binned by difference in coverage and number of mismatches and gaps when aligned to personal assembly and GRCh38. Dashed line represents bins where alignment is “better” (more coverage and/or less mismatches and gaps) to the personal genome than to GRCh38. **c,** Number of lifted off transcripts from personal assembly to GRCh38 by SQANTI structural category with respect to Enhanced GENCODE. Category “other” contains: intergenic, fusion, antisense and genic transcripts. **d,** Percentage of transcripts from personal assemblies over a certain sequence identity threshold with transcripts from GRCh38 based on BLAST. Numbers indicate (left to right) % transcripts with at least 95%, 99%, 99.9% and 100% sequence identity. **e,** Representation of non-reference regions (regions found in a personal genome but not in the reference genome) and their different types based on the mode of transcript overlap**. f,** Density of transcripts (in transcripts/Mb of DNA sequence) discovered in each genome assembly depending whether the transcript is located in a region shared across genome assemblies or intersecting an assembly-specific region. **g,** Number of discovered transcripts intersecting non-reference regions by their relative location with respect to the region. The median is shown at the bottom of the plot. **h,** Most reads that discover a transcript in personal assemblies also (left) map to and (right) are used to discover transcripts in GRCh38. Mean percentage of transcripts from non-reference regions, binned by their percentage of supporting reads (on the left, for mapping; on the right, for supporting the discovery of a transcript in GRCh38). Error bars show standard deviations.

Next, we performed annotation-free transcript discovery in both GRCh38 and the personal assemblies (Methods). We found a median of 16,703 transcripts in personal assemblies; a number similar to that found in GRCh38 (16,702; Methods). To further investigate the similarity between transcripts discovered using the personal assemblies and GRCh38, we lifted over the transcripts from the personal assemblies to GRCh38 and compared them to our Enhanced GENCODE annotation. 98.4% of personal assembly-discovered genes have a sequence identity of ≥99% in GRCh38 after liftover (Supplementary Fig. S26a). On the basis of intron chain coordinate matching, 84.6% of lifted over transcripts are FSMs or ISMs of transcripts in Enhanced GENCODE (Fig. 6c). Moreover, FSMs and ISMs have the highest levels of expression amongst all structural categories (Supplementary Fig. S27; Methods). The remaining lifted over transcripts (14.4%) are unannotated (NICs and NNC) in Enhanced GENCODE and might represent either truly novel transcripts or incorrect liftovers of annotated transcripts. Consequently, using BLAST^53^, we directly compared the sequences of transcripts annotated in the personal genomes with those of the transcripts discovered using GRCh38. A median of 97.2% (15,789) of the transcripts in the personal genomes have a matching transcript built in GRCh38 with at least 99% identity (Fig. 6d; Supplementary Fig. S26b-c). This means that using personal genome assemblies enables the discovery of a modest number of potentially-novel transcripts per sample that are impossible to discover using GRCh38.

Personal genomes include a median of 156 Mb (∼4.7% GRCh38 size) not found in GRCh38 (henceforth non-reference regions; Supplementary Fig. S28; Methods). These non-reference regions are distributed in fragments with a median length of ∼4kb (Supplementary Fig. S28b) and could thus harbor novel transcripts that could not be discovered when mapping against GRCh38 (Fig. 6e). Of the personal assembly-discovered transcripts, only a median of 317 overlap non-reference regions. These represent very few transcripts considering the total size of non-reference regions. Thus, the transcript density of non-reference regions is <2 transcripts per Mb; much lower than in shared regions (∼6 transcripts/Mb; Fig. 6f). Accordingly, non-reference regions have significantly lower sequence complexity (highly repetitive); a characteristic typical of protein-coding and lncRNA-poor regions^54^ (Wilcoxon’s rank-sum, p-value < 2·10^-16^; Supplementary Fig. S28d). From the 317 non-reference region-intersecting transcripts, a median of 89 are fully-contained inside them and 42 have at least one intersecting exon. The remaining 57% (180) only overlap non-reference regions via their introns (Fig. 6g). Furthermore, many reads that contribute to transcripts overlapping non-reference regions do not map (11.6%) or contribute to transcripts (38.3%) in GRCh38; further demonstrating the potential for novel transcript discovery by using personal genomes (Fig. 6h). Nevertheless, there is little transcriptional evidence in non-reference regions compared to the rest of the genome. Analogous comparisons of personal genomes to the telomere-to-telomere (T2T) reference^54^ yield highly similar results, demonstrating that our findings extend to more complete genome assemblies (Supplementary Fig. S26d-g).

Overall, personal genomes enable slightly better mapping and enable the discovery of some potentially novel transcripts that remain invisible when relying on GRCh38. However, non-reference regions appear gene poor considering their sheer total length.

## Discussion

Previous studies have investigated how genetic differences between human populations impact the transcriptome. They have always relied on short-read sequencing technologies to quantify gene expression using reference annotations built on reference genomes^14,22,23,34,55–61^. However, the reference genome is not representative of the human genetic variation and the transcriptome data employed to create the reference annotations is heavily biased towards individuals from European ancestry^6,7,9^. This may seriously limit the potential impact that genomics research has on human health beyond European populations.

To address this limitation, we generated a long-read RNA-seq (LR-RNA-seq) dataset from a population-diverse cohort. This cohort includes samples profiled in other large-scale studies of genetic diversity^19,20,22,23,62^, including the pangenome project^16^. Thanks to the diverse ancestry composition and full-length nature of this dataset, we show that the reference gene annotations widely-used in human genomic research are European-biased; underrepresenting transcripts from non-European individuals. This European-centric bias in transcript representation raises the question of whether other functional elements are equally ancestry biased in genomic catalogs including enhancer, promoter, transcription factor binding, or chromatin mark annotations^63–69^. Since much of genomics research relies heavily on these annotations, any gaps within them render the missing elements virtually invisible to the broader scientific community. This will become increasingly relevant as GWAS studies start to capture a wider diversity of ancestries^70,71^ and therefore will require ancestry-unbiased genome annotations that properly enable the mechanistic understanding of identified genetic variants. Otherwise, we risk overlooking critical aspects of biology and disease and perpetuating healthcare ancestry inequalities.

A recent effort that focuses on overcoming ancestry biases in genome assemblies has created the pangenome graph which represents genetic variation across many human populations^16^. Indeed, we show that personalizing GRCh38 with the sample’s genotype or using full personal genome assemblies enhances the discovery of transcripts that can not be detected with GRCh38. Moreover, we have likely underestimated the benefits of these approaches, given that we rely on haploid linear genomes. Thus, developing tools for transcript discovery with LR-RNA-seq on pangenome graphs will be crucial to annotate the true extent of transcript diversity among human populations. One remarkable result, however, is that the regions of personal genome assemblies not found in GRCh38 or T2T, while accounting for a significant fraction of the genome (∼5%), appear to be depleted in transcriptional activity. This may not be surprising after all, as if the gene density on these regions were comparable to that of the rest of the genome, they would host thousands of genes (protein coding and lncRNAs). Most transcriptome variation across human populations seems to be confined, therefore, to the genome regions that we all share. This suggests that the largest genomic differences between individuals may have little impact on phenotypic traits.

In summary, our work strongly suggests long-read sequencing of transcriptomes from diverse human populations should be prioritized with the ultimate goal of generating the “human pantranscriptome”: the set of all genes and transcripts found in the human species. Indeed, even though our cohort captures only a small fraction of all human genetic variation and profiles a single cell type, we discover thousands of previously unannotated transcripts by sequencing samples from non-European populations. Therefore, expanding the representation of diverse populations in functional genomics across different biological contexts including tissues, cell types, and developmental stages will further close the ancestry gap in genome annotation^58,61,72,73^. Capturing the full spectrum of transcriptomic diversity, particularly from underrepresented populations, will critically contribute to building more comprehensive and equitable genomic resources to advance both scientific discovery and personalized genomic medicine.

## Data and code availability

All supplementary tables can be found on Zenodo: https://zenodo.org/uploads/14219193. All code can be found on GitHub: https://github.com/Mele-Lab/LR-RNA-seq_GeneAnnotationBias.git

Basecalled data and preprocessed FASTQs (>Q10) can be found in Array Express: https://www.ebi.ac.uk/biostudies/arrayexpress/studies/E-MTAB-14935

## Acknowledgements

This project was supported by Award Number 2 U24 HG007234 and U24 HG011451 from the National Human Genome Research Institute, Wellcome Trust [WT222155/Z/20/Z], and the European Molecular Biology Laboratory. We acknowledge support from the Spanish Ministry of Science and Innovation through the Centro de Excelencia Severo Ochoa (CEX2020-001049-S, MCIN/AEI /10.13039/501100011033), the Generalitat de Catalunya through the CERCA programme, and the EMBL partnership. P.C.R. was supported by a predoctoral fellowship “Formación Personal Investigador (FPI)” from “el Ministerio de Ciencia, Innovación y Universidades (MCIN)” and “la Agencia Estatal de Investigación (AEI)” (MCIN/AEI FPI with code PREP2022-000663). F.R. was supported by the fellowship within the “Generación D” initiative, Red.es, Ministerio para la Transformación Digital y de la Función Pública, for talent atraction (C005/24-ED CV1). Funded by the European Union NextGenerationEU funds, through PRTR. M.M. was supported by a grant PID2019-107937GA-I00 funded by MCIN/AEI/10.13039/501100011033 and a grant RYC-2017-22249 funded by MCIN/AEI/10.13039/501100011033 and by “ESF Investing in your future”.

We are grateful to the Guigó laboratory for their valuable input and help with sample handling, as well as to R. Carbonell Garcia from Centre de Regulació Genòmica (CRG) for administrative support. We sincerely thank Tamara Perteghella and Gazaldeep Kaur from CRG for their insights in gene annotation building and filtering and recommendations about the data preprocessing pipeline. Emilio Palumbo from CRG for his advice on the data preprocessing pipeline and his support by managing ONT sequencing software and data transfer. Ferran Reverter and Miquel Calvo from Universitat de Barcelona for their statistics advice. Rebekah Loving from Caltech on her guidance on running lr-kallisto. Samuel Morabito from Centre Nacional d’Anàlisi Genòmica (CNAG) for his general helpful conversations. We thank the Melé laboratory from the Barcelona Supercomputing Center (BSC) for their feedback and recommendations on many analyses. Santiago Marco-Sola from BSC and Álvaro Lucas Barceló from CRG for their insights about the pangenome. **Disclaimer:** The content is solely the responsibility of the authors and does not necessarily represent the official views of the National Human Genome Research Institute or the National Institutes of Health.

## Online methods

### Samples and terminology description

We selected 45 lymphoblastoid cell lines (LCL) from the Coriell Institute for Medical Research biobank from eight different human populations (Supplementary Table S1). Five populations belong fully to the 1000 Genomes Project cohort^19^: 5 Central and Northern European in Utah (CEU), 5 Indian Telugu in the UK (ITU), 6 Peruvian in Lima (PEL), 5 Luhya in Webuye, Kenya (LWK) and 7 Yoruban in Ibadan, Nigeria (YRI). In addition, we selected 3 Han Chinese in Beijing and 1 Han Chinese South from the 1KGP phase 3 cohort, 1 Han Chinese in Beijing from the 1KGP pilot and 1 Han Chinese in Beijing from the HapMap project that we grouped under the label Han Chinese (HAC). Our cohort was completed with 5 Mbuti Pygmy in Congo that we labeled as (MPC) and 6 Ashkenazi Jewish Israeli Population named as (AJI) from the NIGMS Human Genetic Cell Repository Human Variation panel.

Populations were chosen based on the 1000G^19^ results regarding intracontinental diversity based on the number of singletons, balance between genetic variation and population isolation and low levels of admixture with other superpopulations. Individuals were selected to balance sexes and include samples that overlap those in the Human Pangenome Draft^16^.

This study refers to groups of individuals with similar genetics and shared recent evolutionary history, as “populations” or “ancestries” interchangeably. Labels for these groups were adapted from the 1000 Genomes or the Coriell Institute for Medical Research cell lines descriptions, with Fig. 1 showing one of the geographical locations where members of each group are predominantly found today rather than original sampling locations^74^. These labels are not social or ethnic identifiers and have limitations, including the oversimplification of continuous genetic diversity and the admixture present in modern human populations^74^. Accordingly, labels like European, non-European, African and out-of-African refer to genetic descent of individuals and not to the continent where they were born.

### Ethical Statement

This study utilized commercially sourced human cell samples, for which institutional review board approval was not required. All experimental procedures were conducted in accordance with ethical guidelines and principles governing research involving commercially obtained human products.

### Cell Culture and RNA Extraction

Cells sourced from the Coriell Institute were cultured in RPMI complete medium, supplemented with 15% heat-inactivated fetal bovine serum (FBS) and 1% penicillin-streptomycin solution (final concentrations: 100 U/mL penicillin and 100 µg/mL streptomycin). Total RNA was extracted from cell pellets containing 10 million cells using TRIzol reagent, followed by purification with the PureLink RNA Mini Kit (Thermo Fisher Scientific). The purity of the extracted RNA was evaluated using a NanoDrop One spectrophotometer (Thermo Fisher Scientific), and RNA concentration was measured using the Qubit High Sensitivity RNA Assay Kit (Thermo Fisher Scientific, Cat. No. Q32852). RNA integrity and quality were assessed with the RNA High Sensitivity ScreenTape assay on an Agilent 2200 TapeStation system, and RNA Integrity Number (RIN) values were used to confirm sample quality.

### CapTrap, Long-Read Library Preparation and Sequencing Protocol

Total RNA extracted from human samples was used to generate double-stranded cDNA libraries following the CapTrap-Seq protocol^21^, which captures both the 5’ 7-methylguanylate cap and 3’ poly-A tail. A mixture of ERCC and Lexogen SIRV spike-in controls, derived from the SIRV-set 4 (Lexogen), was used in all samples. The spike-in mixture was capped as described in the CapTrap-Seq protocol^21^ and added to the RNA samples prior to cDNA library preparation. To avoid demultiplexing issues, a dedicated flow cell was used for each sample. Resequencing and pooling of reads after preprocessing was performed when a single run did not reach a minimum amount of reads. Sequencing libraries specific to the ONT platform were prepared using the Amplicon by Ligation Kit (SQK-LSK114). Sequencing was performed on a GridION device using ONT R10.4 flow cells, following the standard MinKnow (software version 23.11.7) protocol for ONT’s 114 chemistry.

### Gene annotations

We used GENCODE v47^13^ comprehensive gene annotation downloaded from https://ftp.ebi.ac.uk/pub/databases/gencode/Gencode_human/release_47/gencode.v47.primary_assembly.annotation.gtf.gz and we performed several validations on RefSeq v110 downloaded from https://ftp.ncbi.nlm.nih.gov/genomes/all/annotation_releases/9606/110/GCF_000001405.40_GRCh38.p14/GCF_000001405.40_GRCh38.p14_genomic.gtf.gz. RefSeq contig names were replaced with UCSC format contig names to match those used in GENCODE and enable comparisons. The Enhanced GENCODE annotation was built by adding PODER novel transcripts to GENCODE where the start and end coordinates for gene features were extended to include all novel transcripts (Supplementary Table S7).

### Raw data preprocessing

We basecalled the ONT raw data in Nvidia Hopper GPUs with CUDA (v12.2) using ONT’s dorado tool (v0.5.3) [https://github.com/nanoporetech/dorado] in duplex mode with DNA models dna_r10.4.1_e8.2_400bps_sup@v4.3.0 and dna_r10.4.1_e8.2_5khz_stereo@v1.2. Then we processed the basecalled data in a Snakemake pipeline that removes the parent simplex reads, splits the artificially concatenated reads, also known as chimeric reads, removes the PCR duplicates, cuts all the adapters and filters reads based on mean base quality 7 and 10 (generates both outputs).

We split the chimeric reads using duplex-tools (v0.3.3) [https://github.com/nanoporetech/duplex-tools] with `split_on_adapter` looking for our set of CapTrap-seq adapters containing: ONT sequencing adapters, 5’ and 3’ CapTrap-seq linkers, CapTrap-seq primers and their combinations. Then, all the reads went through a second round of splitting to detect further chimeric reads formed by more than two molecules. All the reads that underwent multiple splittings were removed.

Then we deduplicated by leveraging the 16 nucleotide-long unique molecular identifier (UMI) included inside the 5’ CapTrap-seq linker. ONT sequencing includes random noise at the ends, high error rate, including indel errors, making the UMI position highly variable with respect to the read start. Thus, to find the UMI we used blastn^53^ (blast v2.11.0) to align all reads to a 5’ CapTrap-seq linker database, where the variable UMI corresponds to 16 N. blastn chosen parameters were `-outfmt 5-strand ‘both’ -gapopen 2 -gapextend 2 - penalty-3 - reward 2`. Then we extracted each UMI sequence from the reads by parsing the CIGAR string where the UMI is the sequence corresponding to a 16 nt mismatch. Once the UMI was identified we appended it to the read ID and mapped to the transcriptome (gencode.transcripts.fa) using minimap2^75^ (v2.28) with parameters: `-ax splice --junc-bed` (see Mappings section) `--MD`. After filtering unmapped and supplementary reads, we removed PCR duplicates by running UMI-tools^76^ (v1.0.0) dedup with parameters `--extract-umi-method read_id --method adjacency --edit-distance-threshold 2 --per-contig --per-gene --gene-transcript-map`. This method considers two reads as technical duplicates if they map to the same gene and have concordant UMI with a Hamming edit distance threshold of 2. Overall we could deduplicate a median of 60% of the reads per sample. For the rest of the reads, we could not either identify the 5’ linker, extract a UMI of exactly 16 nt, or they did not map to any annotated transcript. To avoid losing a median of 40% of the reads per sample we assumed that this duplication-unassessed fraction was duplication-free for the rest of the analyses.

After the deduplication we cut all the adapters with porechop^77^ (v0.2.4) with parameters `--min_split_read_size 200 --extra_middle_trim_good_side 2 --extra_middle_trim_bad_side 2`. At this stage, when chimeric reads have already been split, porechop splitting feature removes random noise (<200nt sequences) before or after the adapters surrounding a cDNA read. Finally, reads are filtered for average Q ≥ 7 and Q ≥ 10.

### Mapping

We mapped the preprocessed reads with average Q10 or higher using minimap2^75^ (v2.28) with parameters: `-ax splice --secondary=no --MD -L`. We used a minimap2 generated index with parameter `-x map-ont` created by concatenating GRCh38 primary assembly with Lexogen SIRV-Set 4 FASTA. The input for `--junc-bed` parameter was created from GENCODE comprehensive annotation of primary assembly GRCh38 (from now on referred to GENCODE) concatenated with the Lexogen SIRV-Set 4 annotation using `k8 paftools.js gff2bed` as recommended in minimap2 documentation. Afterwards, we only kept reads mapped to the GRCh38 chromosomes and scaffolds (to discard SIRV reads and ONT library preparation lambda phage spike-ins) and filtered out non-primary alignments, supplementary alignments and unmapped reads.

### Transcript discovery and downsampling

FLAIR, IsoQuant, and ESPRESSO were run using GENCODE v47. We ran FLAIR^25^ (v2.0.0) in two steps: firstly FLAIR correct with default parameters except `--ss_window 8` using GENCODE. Secondly, FLAIR collapse with the same settings as in LRGASP^11^: `--stringent --check_splice --generate_map --annotation_reliant generate`. IsoQuant^26^ (v3.4.1) was run with `--data_type ont--model_construction_strategy sensitive_ont --splice_correction_strategy assembly --delta 8 --genedb GENCODE v47` (or `RefSeq v110`) `--complete_genedb.` ESPRESSO^24^ (v1.4.0) was run with default parameters. LyRic [https://github.com/guigolab/LyRic] (v1.0.4) was executed with the same settings as in the LRGASP project, specifically with `minimumTmergeReadSupport` set to 2 and `exonOverhangTolerance` set to 8. CAGE, DHS signals, and short-read data were not used in the Lyric transcript discovery.

To control for sample throughput in some validations, we randomly downsampled all BAM files to the minimum amount of genomic mapped reads (8,657,393) and ran ESPRESSO as described above. Discovered transcripts across samples were merged by intron chain using Cerberus^5^. ESPRESSO was chosen for the downsample because it automatically removes sub-chains of discovered transcripts (preventing confounding accumulation of truncations) and it has a high sensitivity to detect annotated transcripts and has been benchmarked where it demonstrates good overall performance^24^.

### Transcriptome merging and post-processing

We merged all the transcript models obtained across samples and discovery tools (FLAIR, LyRic, IsoQuant, ESPRESSO) by intron chain keeping the longest transcript start site and end site for each unique intron chain using Cerberus^5^. In brief, we extracted intron chain coordinates using Cerberus and merged those with identical chromosome, intron chain coordinates, and strands across the assemblies; keeping the longest transcript start site and end site for each unique intron chain; a strategy which is somewhat consistent with the GENCODE strategy^78^. Monoexonic transcripts were discarded.

We ran `SQANTI3_qc` from SQANTI3^50^ (v5.2.1) with default parameters on our merged annotation to obtain transcript characteristics like structural category with respect to GENCODE, splice junctions, and associated annotated gene descriptors. Afterwards we assigned each transcript to a gene. We used SQANTI assignments to known genes. However, intergenic transcripts come from novel gene loci, thus we ran buildLoci [https://github.com/julienlag/buildLoci] to group them into genes using default settings while enforcing that novel gene loci only belong to a single strand. We added corresponding gene feature entries in our GTF (using SQANTI assignments for known genes and buildLoci assignments for novel genes). The final annotation from this step makes up the unfiltered, merged PODER annotation (UMA; Supplementary Table S2).

### Transcriptome filtering

We quantified transcript expression using the unfiltered PODER annotation using FLAIR^25^ with `--stringent` parameter. For this step in particular, we used FLAIR to quantify because it does not require gene locus in the GTF file and allows for unambiguous quantification; providing integer counts for how frequently each transcript is seen in the data rather than splitting read counts between transcripts with overlapping structure such as pseudoalignment-based methods such as lr-kallisto^79^. For filtering, this was particularly important to obtain the clearest picture of exactly which intron chains were present in our reads and at what levels.

We built a master table containing all transcripts, associated genes, splice junction information, transcript expression, sample sharing, tool sharing, and SQANTI3 QC descriptors. We additionally added Recount3^80^ splice junction counts data. We downloaded Recount3 splice junctions information from https://snaptron.cs.jhu.edu/data/srav3h/junctions.bgz and kept information for those splice junctions having more than 10 counts. Then we added to our master table the number of samples where a junction has been found and the total sum of counts of the least supported splice junction and the least supported novel splice junction per transcript in our merged annotation (Supplementary Table S3).

Then we filtered our merged gene annotation to generate a high-confidence transcript set. We removed monoexonic transcripts and transcripts associated to genes with biotypeds different from protein-coding and lncRNA. The subsequent filters are based on the properties of transcripts already annotated in GENCODE and on their intron chain novelty as assessed using SQANTI structural categories (Fig. 2b). We applied different filtering criteria depending on the structural classification. For transcript models belonging to annotated genes, we only kept those belonging to protein coding and lncRNA genes because they are 5’ capped and polyadenylated, which are the RNA features that the CapTrap^21^ method selects for. All FSMs were kept. ISMs represented a potential source of 5’/3’ errors, mainly coming from truncations of transcripts during library preparation or basecalling, so consistently with our intron-chain centered approach, all ISMs with a matching FSM were removed. For some transcripts, an ISM was detected but the FSM of the annotated transcript was not. In these cases, the corresponding annotated transcript from GENCODE was added to the annotation to avoid potential loss of read counts when quantifying gene expression. NICs were kept if independently discovered in >2 samples, if expressed with >3 counts in at least 1 sample, and if longer than 300 nucleotides. NNC, intergenic, fusion, genic and antisense transcripts were subject to the same filters as NIC, but additionally were required to be discovered by at least 2 tools and have a minimum Recount3 support ≥25 counts in all splice junctions (Fig. 2b; Supplementary Fig. S4-6). After filtering, we created a GTF with added gene feature entries matching the furthest boundaries of all the transcripts belonging to a gene (Supplementary Table S4-5).

### PODER exon characterization

SQANTI3 quality control was run as described above. For exon characterization, we limited exons to just internal exons to truly only assess the impact of genetics on splicing, as first and last exons are determined by other cellular processes and machinery, and to avoid confounding effects from our method of choosing the longest-transcript start/end site for each intron chain. Using internal PODER exons only, we performed an interval intersect using PyRanges^81^ (v0.0.129) with internal exons from GENCODE. Exons that completely overlapped with the same coordinates were called known. Exons with any degree of overlap but not perfectly-matching start and stop coordinates were called novel 5’/3’. Finally, exons with no reported overlap were called novel.

### ORF prediction

We used ORFanage^82^ (v1.2.0) to predict an initial set of ORFs for each transcript, using the CDSs from GENCODE as a reference. To pick one representative ORF from each transcript we applied ORFanage on, we used the longest match mode of ORFanage. For any transcripts for which ORFanage did not generate any CDS, we performed non-reference-based ORF calling using CPAT^83^ (v3.0.5). To pick one representative ORF from each transcript we applied CPAT on, we considered three cutoffs (0.725, 0.364, and 0) for coding probability as predicted by CPAT. For each transcript, we chose the longest ORF predicted by CPAT, above the highest available cutoff, breaking ties using coding probability within cutoff bins. ORFs returned by CPAT that did not contain a terminal stop codon were not considered.

### Protein annotation and NMD prediction

After the ORF prediction, we ran SQANTI Protein^84^ (v5.2.1) and BLAST protein^85^ (2.11.0). SQANTI protein was run against the CDS of the same reference used to discover novel transcripts while BLAST was run on the GENCODE v47 amino acid sequences. NMD status was predicted using SQANTI protein via the canonical definition of whether the coding sequence of the transcript potentially undergoing NMD or not finishes more than 50 bases before any downstream splice site.

Predicted protein sequence novelty was determined using exact string matching of predicted amino acid sequences against GENCODE annotated sequences from the same gene. We limited these analyses just to novel transcripts deriving from protein-coding loci as reported by SQANTI in PODER. Transcripts predicted to undergo NMD were labeled as NMD, exact matches were labeled as known, substrings were labeled as truncations, extensions of existing amino acid sequences were labeled as extensions, and the rest were labeled as novel. We note that transcripts could have conflicting labels (for instance NMD and known, arising from annotated CDSs known to undergo NMD; or elongation and truncation, arising from overlapping different annotated amino acid sequences as elongations or truncations separately). In these cases, we labeled the transcript with the most conservative annotation of its novelty (*i.e.,* NMD>Known>Known truncation>Known elongation>Novel).

Although we only report protein prediction results for novel transcripts belonging to protein coding genes in this work, we note that we ran the ORF prediction and protein analysis on all PODER transcripts. All metrics and information related to these results can be found in Supplementary Table S5.

### Comparison to external transcriptome assemblies

The GTEx^12^ and ENCODE4^5^ long-read RNA-seq transcriptome annotations were downloaded from https://storage.googleapis.com/adult-gtex/long-read-data/v9/long-read-RNA-seq/flair_filter_transcripts.gtf.gz and https://zenodo.org/records/10869841/files/cerberus.gtf?downloa=1 respectively. The CHESS^31^ transcriptome annotation was downloaded from https://github.com/chess-genome/chess/releases/download/v.3.1.3/chess3.1.3.GRCh38.gtf.gz. We used Cerberus’^5^ `gtf_to_ics` function with default settings to extract unique intron chains from the GTEx, ENCODE4, CHESS, GENCODE v47, RefSeq v110, and PODER annotations. We merged intron chains based on exact coordinates, chromosome, and strand across the annotations to determine which intron chains were present in each annotation (Supplementary Table S6). We ran SQANTI3_qc with default parameters on the GTEx, ENCODE4, and CHESS annotations to understand their novelty composition with respect to GENCODE.

### Variant characterization by VEP

We subset the 1000G^19,20^ VCF to include only those variants found within genes of GENCODE or PODER gene annotations. Then we predicted variant effects using Variant Effect Predictor (VEP)^32^ (v103) in offline mode with GRCh38 reference genome and using either GENCODE or the enhanced annotation (GENCODE enhanced by PODER) as GTF files. The `--everything` and `--total_length` options were applied to provide comprehensive variant annotations, including the length of cDNA, CDS, and protein positions. To assess variant severity between annotations, VEP was run a second time, considering only the most severe impact of each variant (`--most-severe`). The order of severity associated with each predicted consequence followed the Ensembl nomenclature (https://www.ensembl.org/info/genome/variation/prediction/predicted_data.html#consequences).

### Gene and transcript expression quantification

To quantify transcript expression, we used lr-kallisto^79^ (0.51.1) and kb-python^86^ (0.28.2). We ran `kallisto bus` with options `--long --threshold 0.8 -x bulk`; `bustools count` with options `--cm -m`; and `kallisto quant-tcc` with options `--long -P ONT`. Transcript counts were summed to get gene expression (Supplementary Table S11-12).

### Population-specific discovered transcripts and splice junctions

Population-specific transcripts and splice junctions were identified from PODER based on the samples they are discovered in. A transcript or a splice junction discovered in at least two samples (corresponding to between 33% and 50% of all samples from a population) in just one single population was considered population-specific. To test if population-specific features (either transcripts or splice junctions) were enriched in novel categories, we applied a Fisher test per population using all the discovered features in that population as background. To test if European populations have a higher proportion of known splice junctions in population-specific splice-junctions than expected by random chance, we performed random permutation by shuffling population labels across samples, and reassessing population-specific splice junctions. Then we computed the mean proportion of annotated splice junctions by European/non-European groups for every permutation. We ran the permutation test 600 times and calculated the empirical p-value as the proportion of instances having a higher rate of known splice junctions usage than in the original data.

### Tau-based population-specific expressed transcript calling (both from PODER and MAGE)

To call population-specific transcripts on the basis of expression, we used the τ metric of transcript specificity^33^. This indicator provides, for each transcript, a score between 0 (transcript expressed at the same level in all populations) and 1 (transcript expressed in exactly one population) and is computed following the formula:

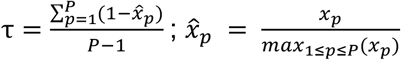

with *x*_p_ being the normalized expression of the transcript of interest in the population *p* and among the *P* populations.

Starting from transcript CPM quantifications from kallisto, we filtered for transcripts expressed ≥1 CPM in ≥1 sample and computed the mean transcript expression per population across the constituent samples. We log-normalized the CPM values and added a pseudocount of 1 before computing τ. Based on the resultant distribution of τ values we called population-specific expressed transcripts as those with τ ≥ 0.8.

For the PODER results, we used the quantification transcript-level quantification values from the LR-RNA-seq data quantified using the PODER annotation with lr-kallisto (Supplementary Table S5, S11). For the MAGE results, we used Enhanced GENCODE to quantify transcript expression using kallisto (Supplementary Table S10, S13).

### MAGE data processing

We downloaded all FASTQs from the MAGE dataset^23^ from the European Nucleotide Archive: https://www.ebi.ac.uk/ena/browser/view/PRJNA851328. Data were quantified using kallisto^87^ (0.51.1) using the options `-x BULK --parity=paired --stran=unstranded --tcc--matrix-to-directories`. We quantified the data twice, once using GENCODE v47 as the transcriptome reference, and once using Enhanced GENCODE (GENCODE v47 + novel PODER transcripts) as input (Supplementary Table S8-10). We computed the number of reads quantified in each case by summing up the counts in each output matrix and computed the ratio between the number of reads quantified using each annotation.

### Allele specific analysis

We downloaded the high-coverage 1000G^20^ imputed and phased VCF from http://ftp.1000genomes.ebi.ac.uk/vol1/ftp/data_collections/1000G_2504_high_coverage/working/20220422_3202_phased_SNV_INDEL_SV/ and subsetted the 30 samples overlapping our dataset. Then, we processed it following LORALS^12^ recommended data preprocessing. We ran allele-specific analysis using LORALS with default parameters on our mapped Q7 reads and with all the optional filterings for problematic regions activated as described in the documentation [https://github.com/LappalainenLab/lorals]. As part of the analysis, LORALS generates two reference genomes for each sample that represent the sample’s phased, biallelic SNP haplotypes in GRCh38. We only kept ASE results with more than 20 counts per gene and calculated FDR after all filterings. ASTU was computed by running a chi-squared test on those genes that have at least two transcripts with at least 10 counts and have counts on both alleles (Supplementary Table S14-15).

### GWAS hit enrichments

ASE and ASTU significant genes were tested for enrichment in GWAS catalog^39^. We downloaded the GWAS catalog “All associations v1.0” and for each trait extracted SNP_GENE_IDS if the GWAS hit was genic and UPSTREAM_GENE_ID and DOWNSTREAM_GENE_ID if it was intergenic, which contain the ENSEMBL gene ID of the genes included in the column MAPPED_GENE. Then we ran Fisher’s test using this GWAS trait built gene sets and ASE and ASTU tested genes as background. We set a significance threshold at FDR = 0.02 (Supplementary Table S16).

### Personalized-GRCh38 analysis

Personalized versions of GRCh38 were created for all biallelic SNPs from the 1000G^19,20^ VCF for the 30 overlapping samples using LORALS^12^. Two personalized versions were created for each sample; representing the two phased haplotypes. LR-RNA-seq Q10 FASTQ reads were mapped to each resultant haplotype and to GRCh38 with minimap2^75^ (v2.28) and parameters `-ax splice --MD --secondary=no -L`. We removed unmapping, secondary, and supplementary alignments. Then we ran ESPRESSO^24^ as described above, using the same genome (either personalized-GRCh38 haplotype or GRCh38 to perform transcript discovery. We ran SQANTI^50^ using previously-specified options to extract the coordinates and novelty of each splice junction and transcript with respect to GENCODE. We used Cerberus^5^ to extract the coordinates of each intron chain. We used exact genomic coordinate matching for intron chains and splice junctions to determine which features were detected for each sample for the three different reference genomes (Supplementary Table S17).

For the analyses related to novel transcripts, we specifically only used transcripts with novel splicing (*i.e.,* we removed all transcripts that FSMs and ISMs), as we expect that gains in novelty are specifically related to allowing novel splicing events rather than truncations of existing splice chains.

To find the variants from each sample that intersect with the splice junction-proximal exonic regions and splice sites, we used the splice junctions from SQANTI to obtain 1) ±10bp exonic regions and 2) 2bp intronic splice site regions. We intersected the regions with the 1000G VCF^19,20^ for the corresponding sample, and recorded, for each splice junction, whether an alternate biallelic SNP was present.

### Computing FSTs in internal exons

Weir and Cockerham Fixation Index (F_ST_) was computed using VCFtools (v0.1.16) (`--weir-fst-pop` option) and for each pair of populations covering our 30 samples genotyped in the 1000G^20^. CHB was used as a proxy for our HAC group. To be able to compare these values, only biallelic SNPs were kept.

We extracted known exons and regions of novel exons that are not in the GENCODE v47 annotation using PyRanges’ implementation of subtract (v0.0.129)^81^. Using the resultant BED regions, we intersected the F_ST_ values with the exons and computed the mean F_ST_ value across variant positions and contrasts (where only considering the 5 contrasts involving CEU, as CEU best represents the ancestry on which the gene annotation is based and biased towards). For statistical testing, we stratified the mean F_ST_ values by Novel and Novel 5’/3’ based on the exon of origin as previously described.

### Personal genome mapping

Assembly comparisons were performed using the six maternal haplotypes from the Human Pangenome Reference Consortium (HPRC) samples overlapping those sequenced in our LR-RNA-seq dataset (HG002, HG00621, HG01928, HG01952, NA18096, NA19240) as well as the two reference genomes, GRCh38 and T2T^54^ respectively. Personal assemblies were downloaded from the pangenome resource (https://s3-us-west-2.amazonaws.com/human-pangenomics/index.html?prefix=working/).

For the read mapping, firstly, each LR-RNA-seq sample was aligned to the two reference genomes as well as to its corresponding personal genome, using minimap2^75^ (v2.28) with the -`ax splice --secondary=no --MD -L`. Then, only primary alignments were considered and used as input in IsoQuant^26^ (v3.6) in annotation-free mode to reconstruct transcript models on each assemblies and produced expression quantification in counts and TPM. Read identifiers were used to track read matches between personal and reference genomes. Comparison of “coverage” reads between assemblies was carried out on the basis of the CIGAR provided through the alignment.

For the genomic comparison, minimap2 (v2.28) with the `-ax asm5` profile was used to compute assembly alignments and the `genomecov` module of BEDtools^88^ (v2.30.0) was applied to identify the unmapping genomic region. All regions smaller than 1000bp were filtered and the sequences associated with these regions were extracted from the original genome using the `getfasta` module. The complexity and repetitiveness of genomic sequences (whole genome or extracted regions), were quantified by calculating the Shannon entropy after counting the number of 6-mers present in each sequence. As a control, for each comparison, the Shannon entropy was computed selecting same size random sequences using the `shuffle` module of BEDtools (v2.30.0) with the `-noOverlapping` option (`-seed 160524`). To establish if a transcript was identified in an unmapped genomic region, the `intersect` module of BEDtools (v2.30.0) was used (with and without option `-F 1.00` to detect inclusion). Three categories of intersection were defined: “Fully included” if the transcript was entirely overlapped by a region, “Only intronic” if the region was entirely overlapped by a transcript intron, “Exonic intersect” if at least 1bp of a transcript exon was intersecting such a region.

### Comparison between Enhanced Gencode annotation and transcripts identified through personal assemblies

Transcripts identified through IsoQuant^26^ (v3.6) in the personal genome were lifted to the GRCh38 reference assembly using the LiftOff pipeline^89^, which is based on the minimap2 mapper (parameters: `-a --end-bonus 5 --eqx -N 50 -p 0.5`). Initially, a database was constructed from the analyzed annotation using the `-g` option, which was subsequently used with the `-db` option. When gene or transcript models were absent in the provided annotation, the `-infer_genes` and `-infer_transcripts` options were applied to infer these models. In addition to the lifted coordinates, two measures are computed: coverage, which represents the proportion of the reference gene or transcript length that matches the target genome; and sequence identity, which measures the percentage of aligned nucleotide positions that remain identical between reference and lifted annotations. Then, `SQANTI3_qc` from SQANTI3^50^ (v5.2.1) was run with the parameters `--force_id_ignore --skipORF` to obtain transcript characteristics and classification according to the merged annotation (Enhanced GENCODE by novel PODER transcripts).

### Comparison between transcripts identified using personal and reference assemblies

For transcripts identified through IsoQuant^26^ (v3.6) using the same sample but mapped to its own personal assembly and to the references ones, the FASTA sequences were extracted using gffread (v0.12.8) `-w` option. Then, a blastn (BLAST v2.12.0+) was performed (option `-outfmt 6`) considering the transcripts discovered when using the personal genome and those discovered when using the GRCh38 reference genome, as query and subject respectively. For each transcript, the best alignment was extracted using the `-max-hsps 1` and -`max-target-seq 1` parameters.

