## Supplementary Materials for "Long-read transcriptomics of a diverse human cohort reveals widespread ancestry bias in gene annotation"

### Supplementary results

##### Transcript catalog overlap of PODER transcripts

Novel transcripts in PODER might still be discovered by other transcript discovery efforts. First, we assessed the overlap and composition of PODER transcripts with large-scale RNA-seq derived transcriptome efforts, including full-length (ENCODE[1] and GTEx[2]) and short-read datasets (CHESS[3]; Supplementary Table S6). Of the four, PODER has the highest proportion of FSM transcripts compared to GENCODE (Supplementary Fig. S10a; Methods). In addition, it has the lowest number of novel intron chains not overlapping other catalogs (~31k) despite having the highest sequencing throughput among the LR-RNA-seq derived resources (Supplementary Fig. S10c). Given the differences in sample composition, data processing strategies, and sequencing throughput, we did not expect high overlap between transcripts. Accordingly, 59.3% of FSMs and 44.8% of NICs are detected in at least one of these catalogs, however NNCs have a lower overlap (10.3%; Fig. 2c; Supplementary Fig. S10b-c). Despite the low overall overlap of novel PODER transcripts (24.7%), it is comparable to the transcript sharing from the other three annotations (20.0-35.9%). Finally, we show that novel, externally-unsupported PODER transcripts are lowly-expressed, suggesting that differences in throughput and detection sensitivity may have played a role in their lack of external reproducibility (Wilcoxon rank-sum, $p<2.22e-16$; Supplementary Fig. S8-10; Supplementary Results). In general, we found that PODER has a comparable overlap with other RNA-seq based transcriptome annotation efforts despite fundamental differences between the catalogs.

##### Gene and transcript quantification using PODER

We used PODER to quantify gene and transcript expression in our dataset to further characterize our cross-ancestry gene annotation. We detected a median of ~16k genes and ~63k transcripts per sample with at least one count, the latter being highly dependent on sequencing depth as also seen in transcript discovery. As expected, most expressed genes in a sample are protein coding (median = 79.2%) and demonstrate high sample sharing. In contrast, the remaining expressed transcripts are either from lncRNA or novel genes with lower sample sharing (Supplementary Fig. S8-9; Supplementary Table S11-12).

##### Comparing predicted variant effects in GENCODE and PODER-enhanced GENCODE

To assess how exonic and splice site variants effects change when adding our novel transcripts, we ran variant effect predictor (VEP; Methods)[4]. We used both GENCODE and a version of GENCODE with novel transcripts from PODER along with their predicted CDSs, which we call enhanced GENCODE. Using the variants present from our sampled populations from the 1000 Genomes[5], we compared the global predicted effects of variants across the three annotations. We found 302,615 novel variant consequences (13.4% increase), although by and large the proportions of variant effect impact are nearly the same between the three annotations and there are very few changes in variant consequence (Supplementary Fig. S13). This suggests that our novel transcripts are functionally comparable to annotated ones and align with evolutionary constraints, reinforcing their plausibility and biological relevance.

##### Validation of transcript discovery bias in reference annotations

We validate the transcript discovery bias, which shows a higher discovery of novel transcripts in non-European populations than in European populations in different ways. First, we check that this effect is not driven by genetic heterogeneity differences between Africans and out-of-Africa populations, which show no significant differences (Supplementary Fig. S14a). Secondly, we observe that the filtering strategy is not causing this effect by performing the comparison on the unfiltered set of transcripts (unfiltered merged annotation or UMA) that was used to generate PODER (Supplementary Fig. S14b). Then, we downsampled to the minimum sequencing depth (~8.6 million reads) and ran ESPRESSO transcript discovery tool[6], guided by either GENCODE or RefSeq, to find out if the sequencing depth normalization was reliable. We found no significant differences in the number of overall discovered transcripts between populations (Supplementary Fig. S14c). Upon stratification by structural category, we observe significantly more novel transcripts (both NIC and NNC) in non-European populations compared to Europeans both when using GENCODE and RefSeq as a reference (Supplementary Fig. S14d-e).

##### Validation of population-specific discovered transcripts

In order to validate our finding that population-specific transcripts discovered in European populations are more frequently annotated, we performed several orthogonal analyses. First, we verified that these effects were not an artifactual product of our filtering strategy by repeating the population-specific transcript discovery analysis with the whole unfiltered merged annotation (UMA; Supplementary Table S2-3), which includes all protein coding and lncRNA transcripts that are multiexonic regardless of whether they passed filtering. With UMA, we see a similar trend where all non-European confidence intervals are higher and almost non overlapping with European ones, thus demonstrating that the filtering is not biasing the results (Supplementary Fig. S16b). To ensure that differences in sequencing depth or number of unique samples per population did not affect this result, we downsampled all populations to 4 samples and 8.6 million reads per sample, and performed transcript discovery with ESPRESSO guided by GENCODE and RefSeq, independently. Again, we observe the same trends (Supplementary Fig. S16c-d, Methods). To overcome uneven limited sample sizes within populations, we grouped populations into European and non-European to define a higher sample sharing threshold (33% sharing; 4 and 11 samples, respectively) to identify transcripts specific to these 2 groups. We found a significant enrichment of FSM in European-specific transcripts with 89.4% of European group-specific transcripts being FSMs (42/47) versus 40.7% (132/324) non-Europeans (Fisher's test, $p=10^{-10}$, OR=12.1). As the novelty of transcripts is defined based on the annotation status of their splice junctions, we further validated our results at the splice junction level as well by calling population-specific splice junctions. We found that population-specific splice junctions from European populations are more often already annotated than expected by random groups of samples (permutation test, empirical p=0.015) but those from non-European populations are not, which demonstrates that this is not a spurious result (permutation test; empirical p=0.88; Methods). Altogether, these results suggest that the overrepresentation of European-specific transcripts in annotations is robust and driven by their novel splice junction content.

**ASTU testability**

For ASTU testing, a gene requires two transcripts with a minimum expression (10 counts). When using PODER, there are more novel expressed transcripts to genes that in GENCODE have a single expressed transcript in our dataset, therefore increasing gene testability (Supplementary Fig. S24; Supplementary Table S15).

### Supplementary tables

**Table S1:** Sample and experimental metadata pertaining to each LR-RNA-seq sample.
**Table S2:** Unfiltered merged annotation (UMA) GTF for all spliced intron chains discovered by each of the tools.
**Table S3:** Metadata table for SQANTI QC, Recount3 support, protein prediction, and other relevant characteristics for each transcript in the UMA annotation used to filter transcripts.
**Table S4:** Final PODER GTF, including predicted CDSs for novel transcripts and annotated CDSs for known transcripts.
**Table S5:** Metadata table for SQANTI QC, Recount3 support, protein prediction, and other relevant characteristics for each transcript in PODER.
**Table S6:** Boolean detection of transcripts across PODER, annotations (GENCODE, RefSeq) and other RNA-seq / LR-RNA-seq-derived transcript catalogs (CHESS, GTEx, ENCODE4).
**Table S7:** Enhanced GENCODE GTF with novel transcripts from PODER added to all annotated transcripts from GENCODE v47.
**Table S8:** Transcript counts for the MAGE RNA-seq dataset computed using the PODER annotation with kallisto.
**Table S9:** Transcript counts for the MAGE RNA-seq dataset computed using the GENCODE annotation with kallisto.
**Table S10:** Transcript counts for the MAGE RNA-seq dataset computed using the Enhanced GENCODE annotation with kallisto.
**Table S11:** PODER transcript counts in each sample as quantified by lr-kallisto.
**Table S12:** PODER gene counts in each sample as quantified by lr-kallisto.
**Table S13:** Tau values for Enhanced GENCODE transcripts computed on the MAGE RNA-seq dataset.
**Table S14:** Allele-specific expression results.
**Table S15:** Allele-specific transcript usage results.

**Table S16:** GWAS enrichment results for ASTU genes overlapping GWAS genes.
**Table S17:** Isoforms and their intron chains detected using personalized-GRCh38s.

### Supplementary figures

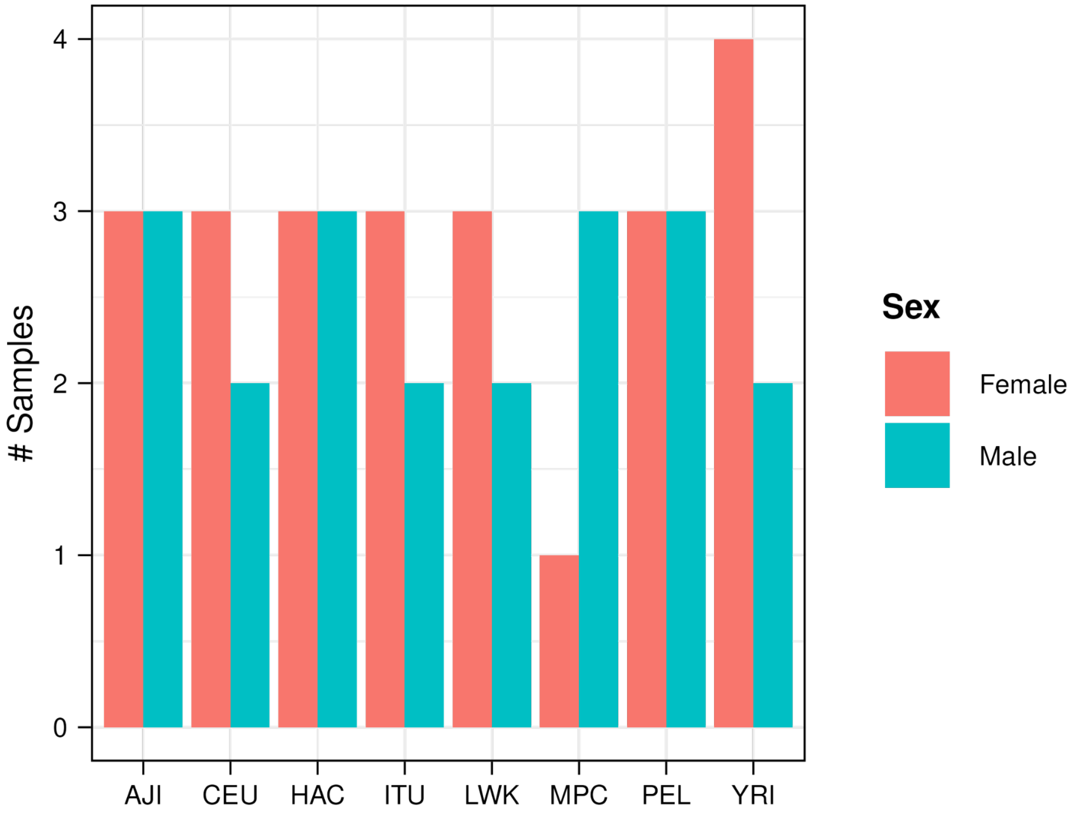


**Supplementary Fig. S1: Samples within most populations are well sex-balanced.**
Number of samples per population by sex.

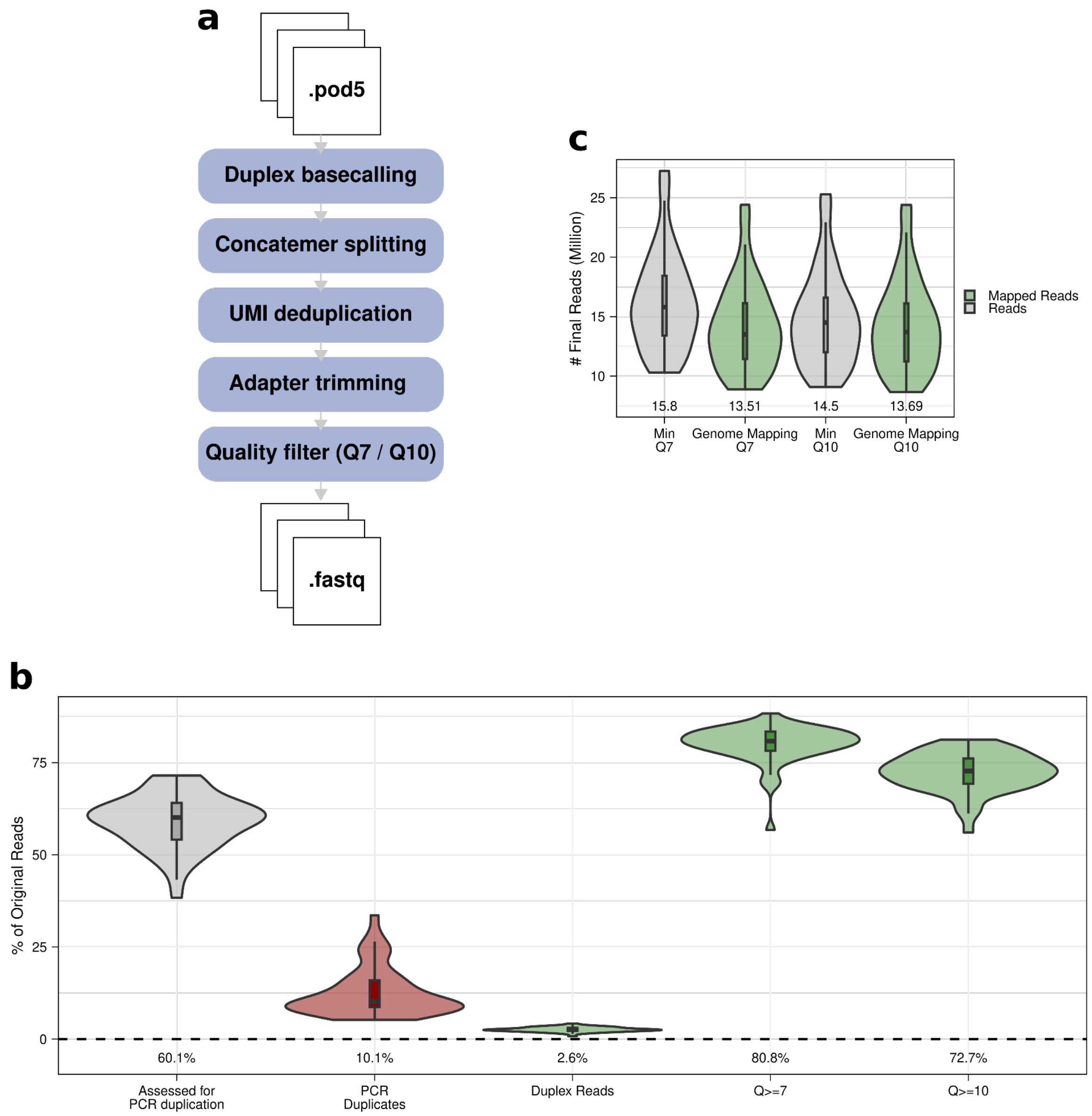


**Supplementary Fig. S2: Oxford Nanopore preprocessing pipeline and number of reads affected by each step.**
**a**, Schema of ONT preprocessing pipeline.
**b,** Percentage of obtained reads involved (grey), removed (red) and kept (green) in each step.
**c,** Final number of reads after the preprocessing pipeline with different minimum thresholds of average read quality (grey) and number of genome mapped reads. Genome mapping Q7: GRCh38 mapped >Q7 reads with minimap2 with GENCODE v47 splice junction guidance. Genome mapping Q10: >Q10 reads; does not have gene annotation splice junction information.

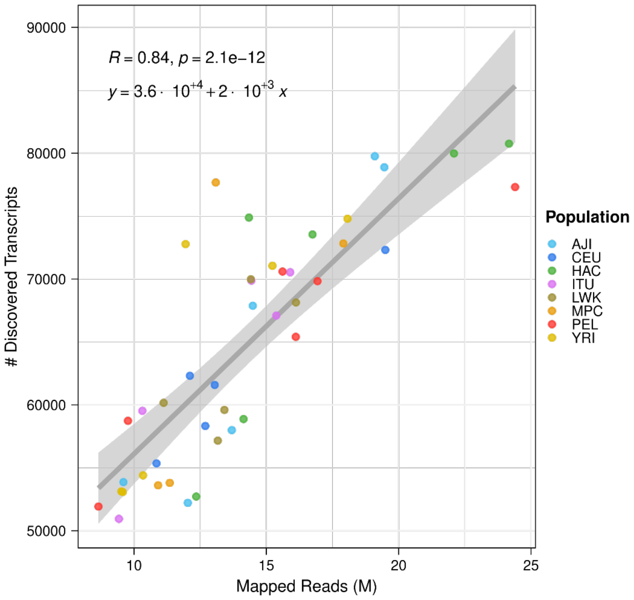


**Supplementary Fig. S3: Transcript discovery is highly dependent on the number of mapped reads.**
Number of discovered transcripts versus the number of million mapped reads per sample (*R*: Pearson's correlation coefficient).

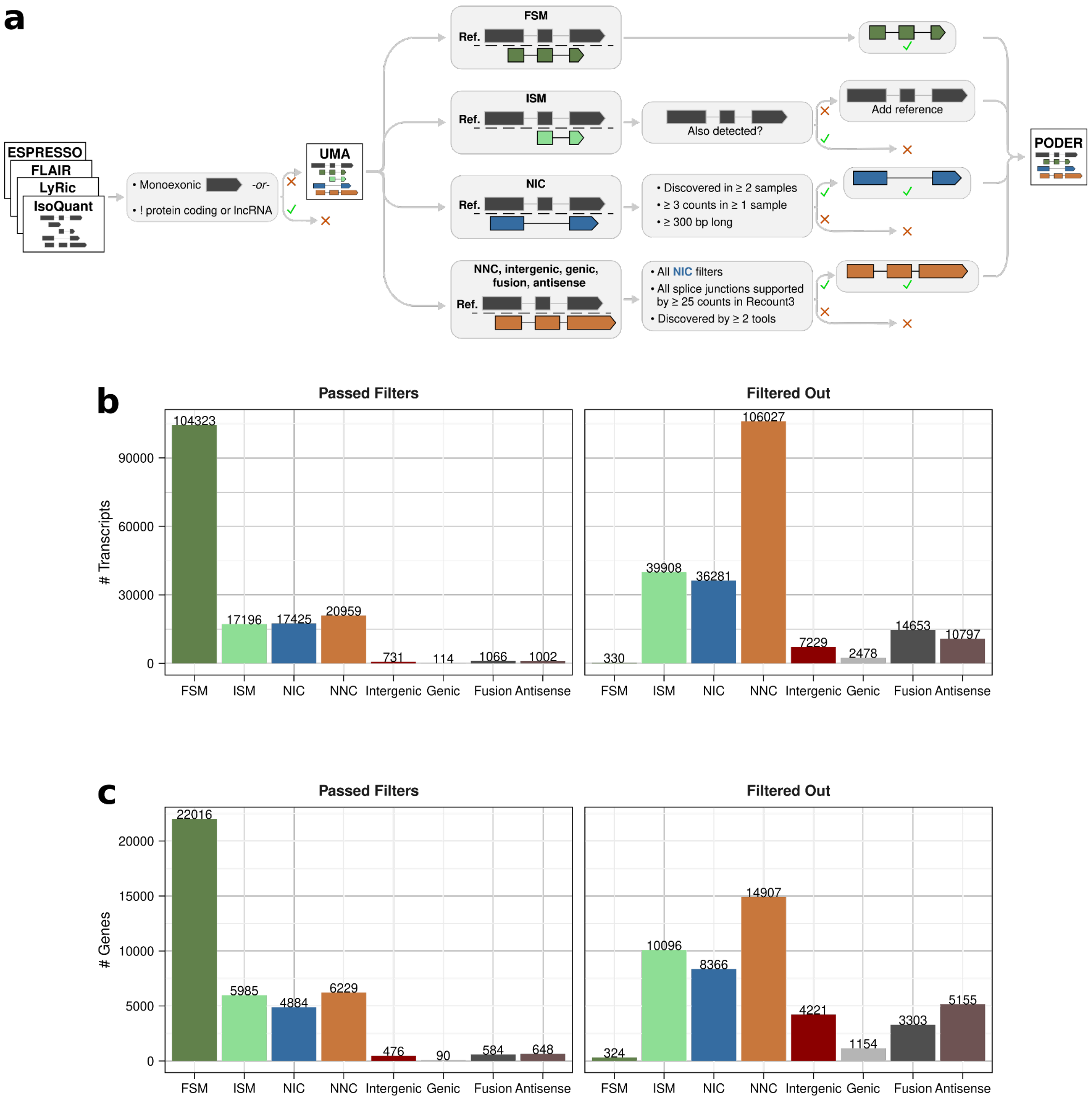


**Supplementary Fig. S4: PODER full-length transcriptome filtering strategy and results overview.**
**a,** Transcript merging and filtering strategies applied to transcripts of different SQANTI structural categories.
**b,** Number of transcripts (left) passing and (right) failing filtering. Note that passing ISMs were all replaced by GENCODE v47 transcripts matching their intron chains in the final annotation. Also note that failing FSM transcripts are from genes whose biotypes are not protein-coding or lncRNA.
**c,** Number of genes to which transcripts (left) passing and (right) failing filtering belong to.

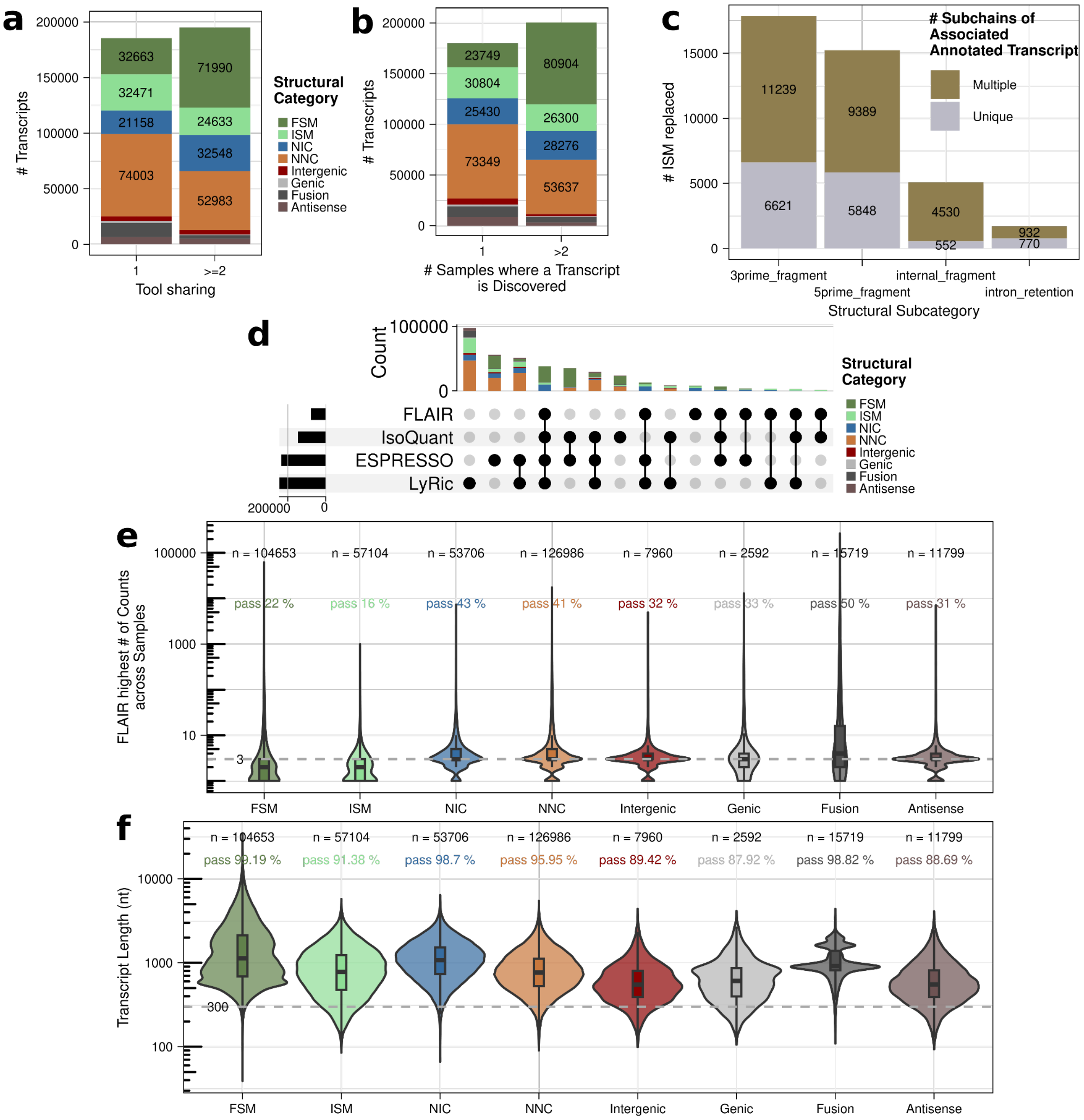


**Supplementary Fig. S5: PODER full-length transcriptome tool, sample, expression, and length-based filtering.**

**a,** Number of unique unfiltered transcripts per structural category based on if they are found by just 1 transcript discovery tool or by ≥2 tools.

**b,** Number of unfiltered transcripts in one or more samples.

**c,** Number of ISMs without a matching FSM discovered in our dataset by structural subcategory, colored by whether they are the unique subchains of a FSM or if there are multiple ISM representing different subchains of the same FSM.

**d,** Intersection of transcripts before filtering discovered by each transcript discovery tool, colored by structural category.

**e,** FLAIR counts (maximum across samples) for transcripts before filtering with length cutoff and number of transcripts passing filtering indicated per structural category.

**f,** Transcript length distributions for transcripts before filtering with length cutoff and number of transcripts passing filtering indicated per structural category.

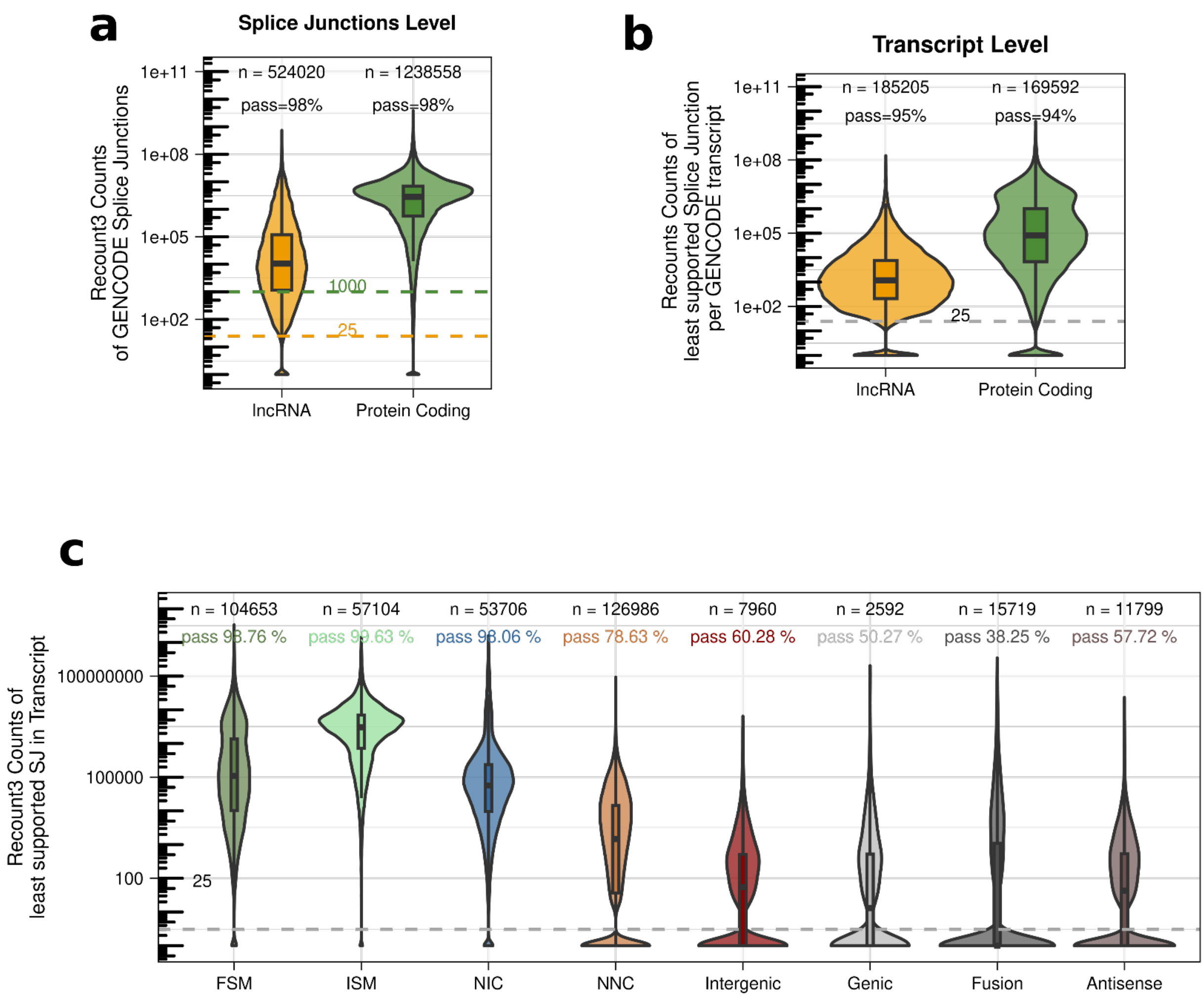


**Supplementary Fig. S6: PODER full-length transcriptome and splice junction support based filtering.**
**a,** Recount3 splice junction support counts for GENCODE splice junctions split by gene biotype.
**b,** Recount3 splice junction support counts for least-supported splice junction per GENCODE transcript, split by gene biotype.
**c,** Recount3 splice junction support counts for least-supported splice junction per transcript from transcripts before filtering with cutoff and PERCENTAGE of transcripts passing filtering indicated per structural category.

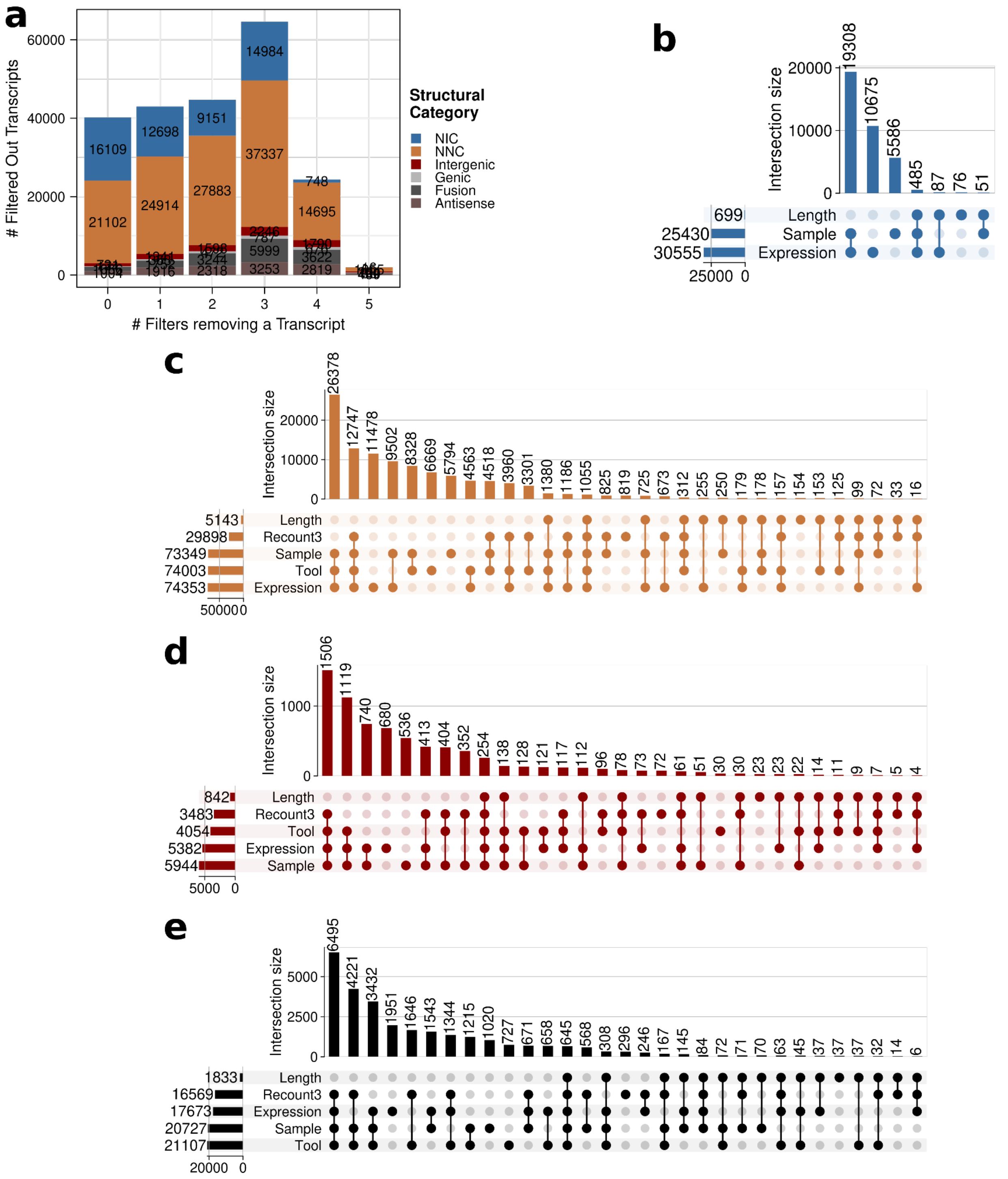


**Supplementary Fig. S7: High redundancy between different filters for novel transcripts.**
**a,** Number of filtered-out transcripts by the number of different filters it was removed by and structural category.
**b-e,** Number of filtered-out transcripts removed by different combinations of filters for **b,** NIC **c,** NNC, **d,** Intergenic, **e,** Antisense, Genic, and Fusion transcripts.

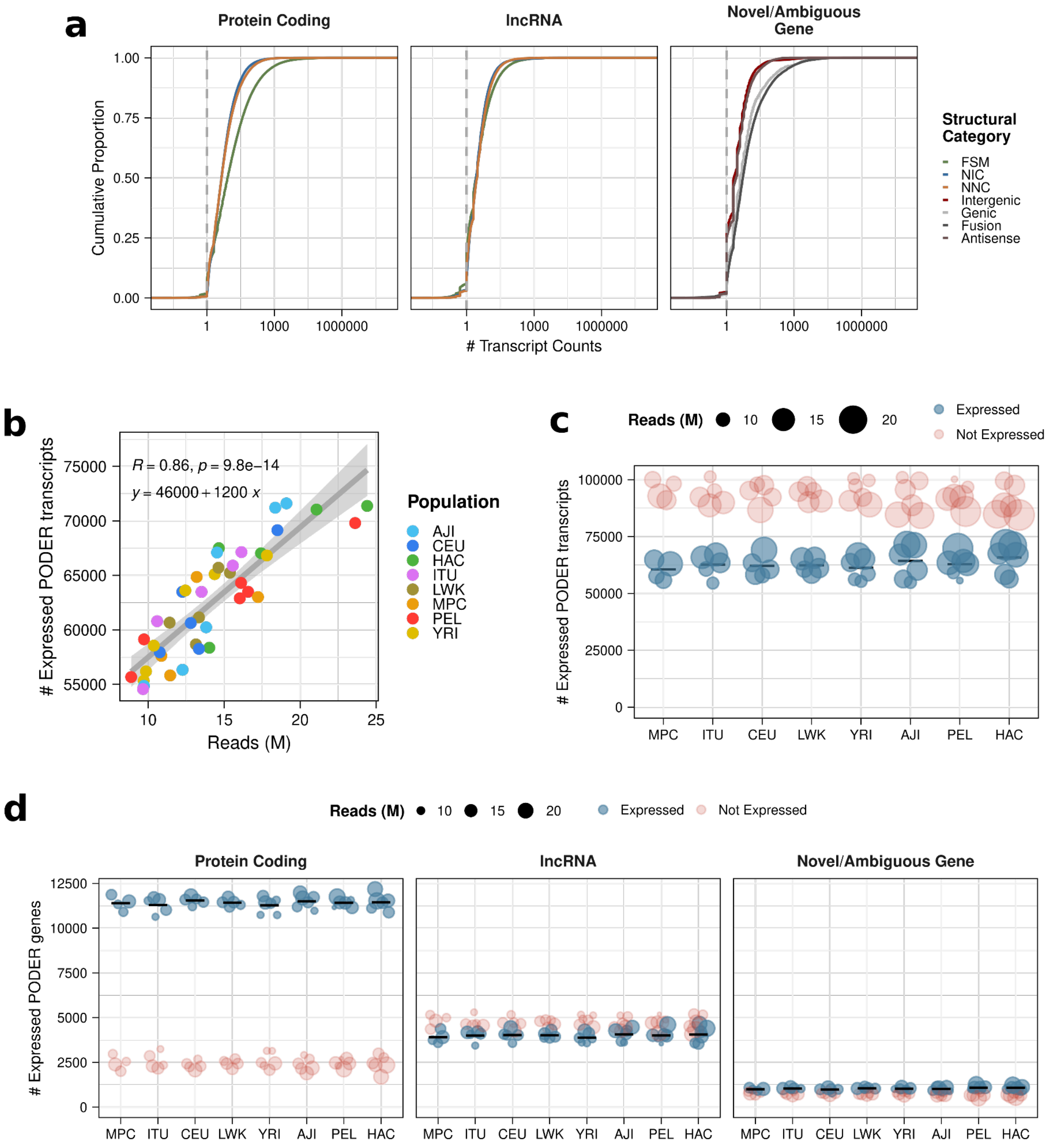


**Supplementary Fig. S8: Gene and transcript expression in PODER.**

**a,** Empirical cumulative distribution plot of PODER transcript expression (counts) stratified by gene biotype and transcript structural category.

**b,** Number of expressed PODER transcripts versus number of reads per sample.

**c,** Number of expressed and unexpressed PODER transcripts per sample stratified by population.

**d,** Number of expressed and unexpressed PODER genes per sample stratified by population and gene biotype.

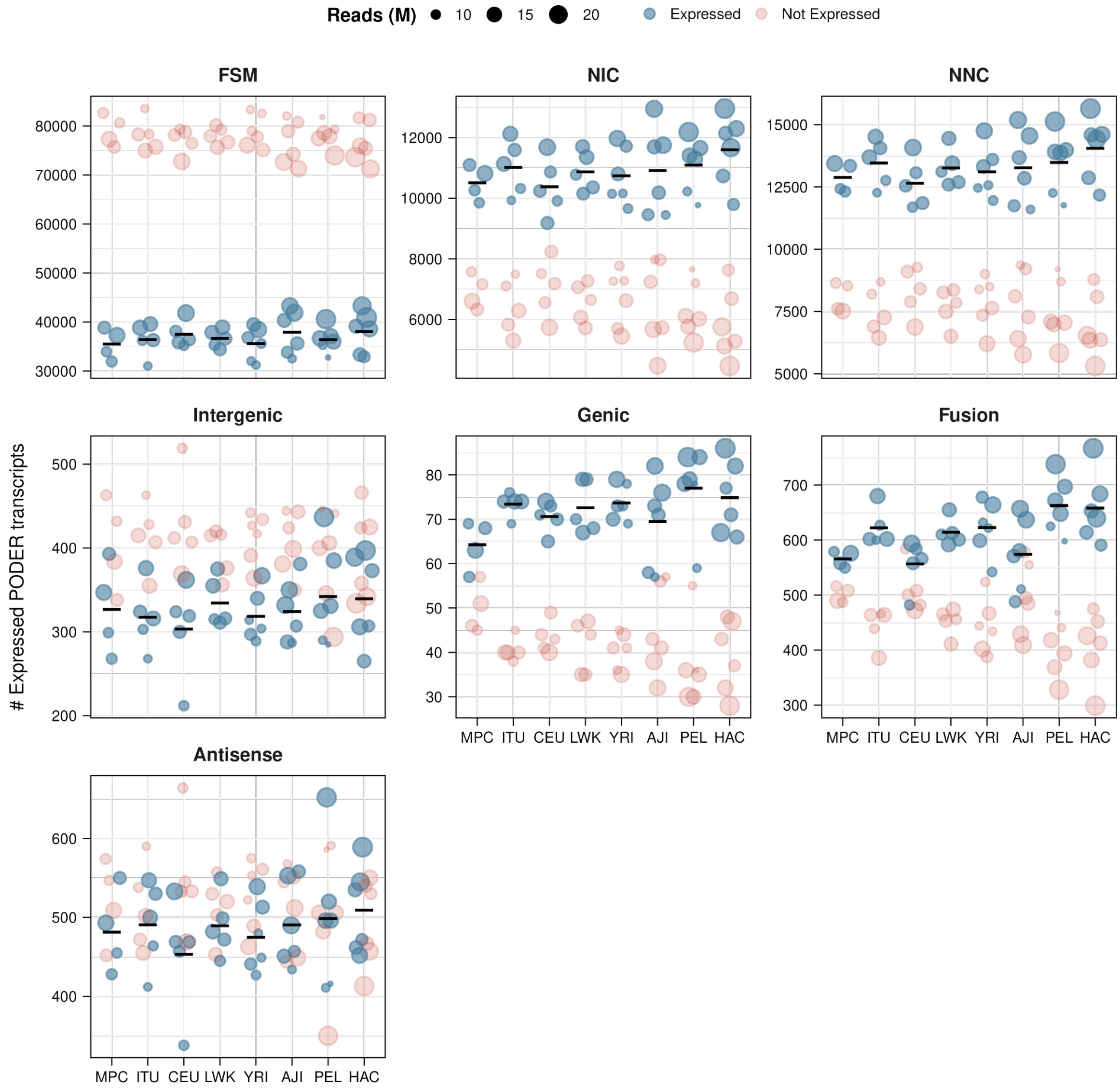


**Supplementary Fig. S9: Transcript expression in PODER.**
Number of expressed and unexpressed PODER transcripts per sample stratified by population and structural category.

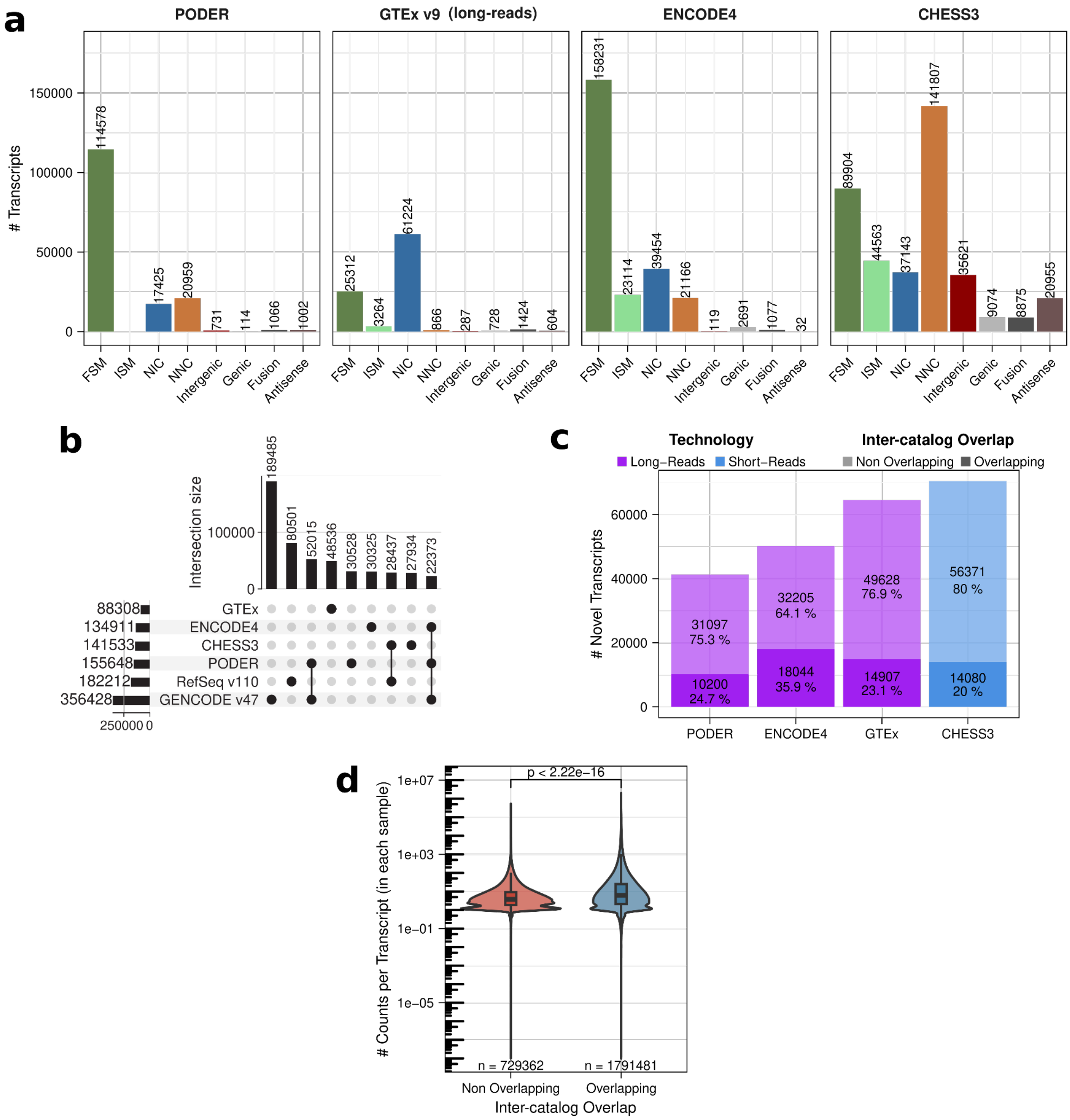


**Supplementary Fig. S10: Support for PODER transcripts beyond GENCODE.**

**a,** Number of transcripts from PODER, GTEx v9, ENCODE4 (LR-RNA-seq derived); and CHESS3 (short-read RNA-seq derived) by structural category with respect to GENCODE v47.

**b,** Intersection and number of intron chains reported across GENCODE v47, RefSeq v110, full-length transcriptome annotation efforts (ENCODE4, GTEx, PODER), and short-read transcriptome annotation efforts (CHESS3). Only the top 9 intersections are shown. Full set size numbers shown for each annotation on the left.

**c,** Number and percent of novel transcripts (with respect to GENCODE) per transcript catalog that overlap or not with any of the others (eg., # and % of novel PODER transcripts that are and are not in ENCODE4, CHESS3, or GTEx).

**d,** Expression values (counts) for novel PODER transcripts, split by whether they are overlapping or not between at least two catalogs (Wilcoxon rank-sum test).

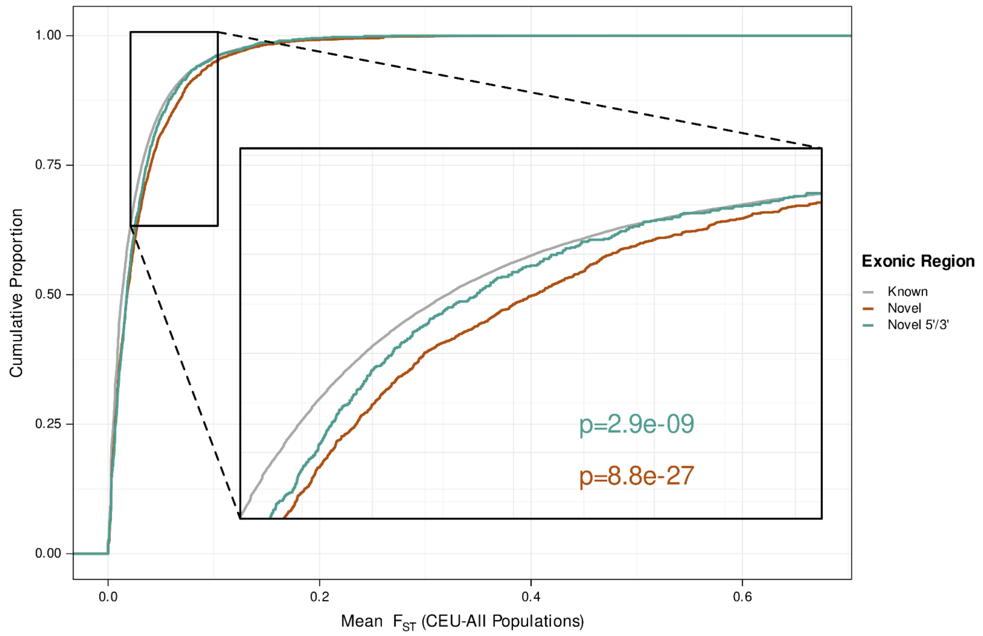


**Supplementary Fig. S11: Genetic differences between populations are related to exon annotation status.**
Cumulative proportion of mean Wright's fixation index ($F_{ST}$) between CEU and YRI, LWK, HAC, PEL, ITU, stratified by exonic regions that are known, novel or overlapping a known exon (novel 5'/3'). $F_{ST}$ is a metric that compares SNPs allele frequencies between two populations; it goes from 0 to 1: the higher the $F_{ST}$ the higher the differences in allele frequencies. Wilcoxon's rank-sum test.

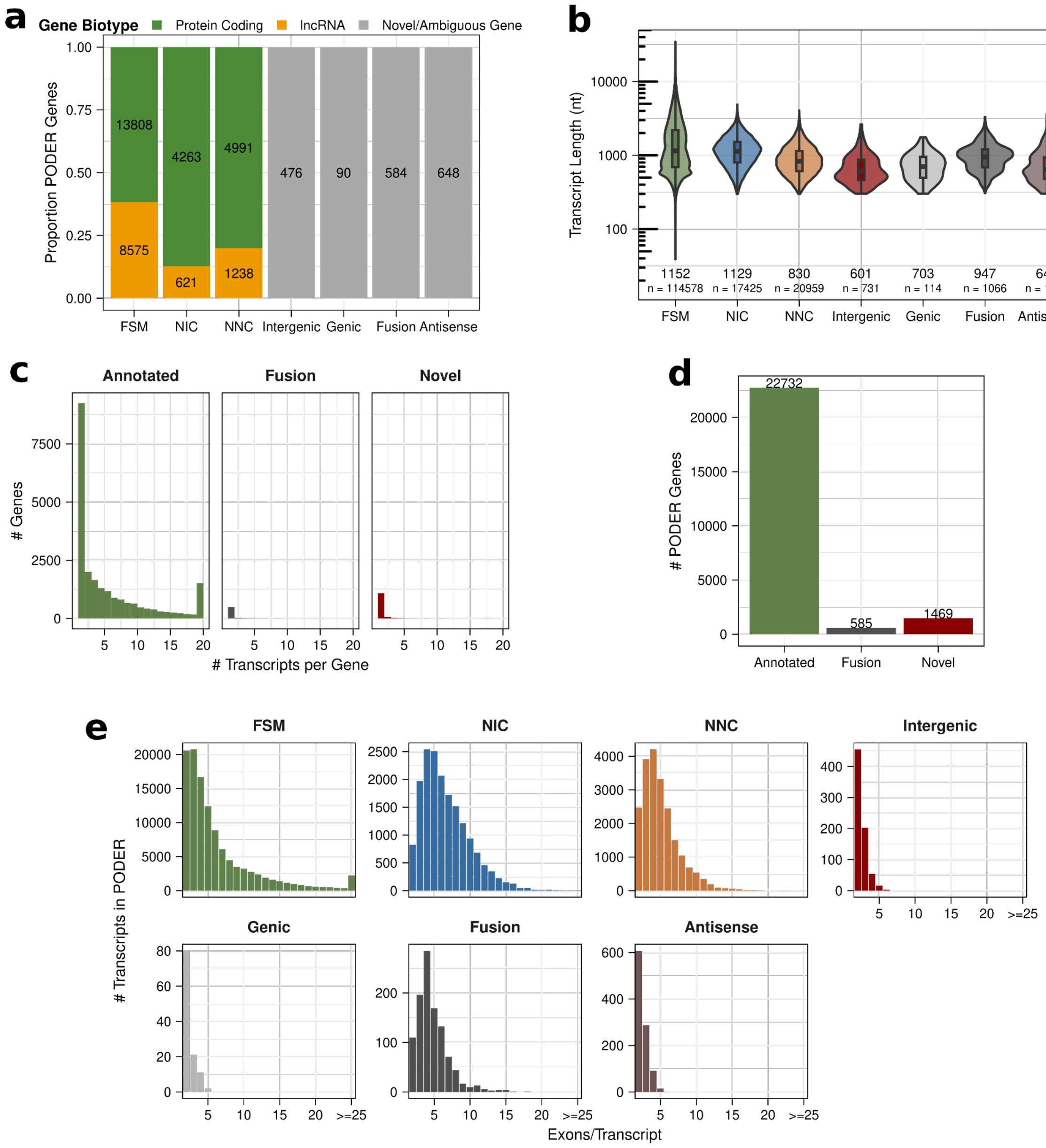


**Supplementary Fig. S12: PODER transcriptome characterization.**

**a,** Number of genes with transcripts in PODER in different structural categories, colored by gene biotype of parent gene.

**b,** Length distributions of PODER transcripts by structural category.

**c,** Number of transcripts in PODER per gene for known, fusion and novel genes.

**d,** Total number of known and novel genes in PODER.

**e,** Histogram of number of exons per transcript in PODER split by structural category.

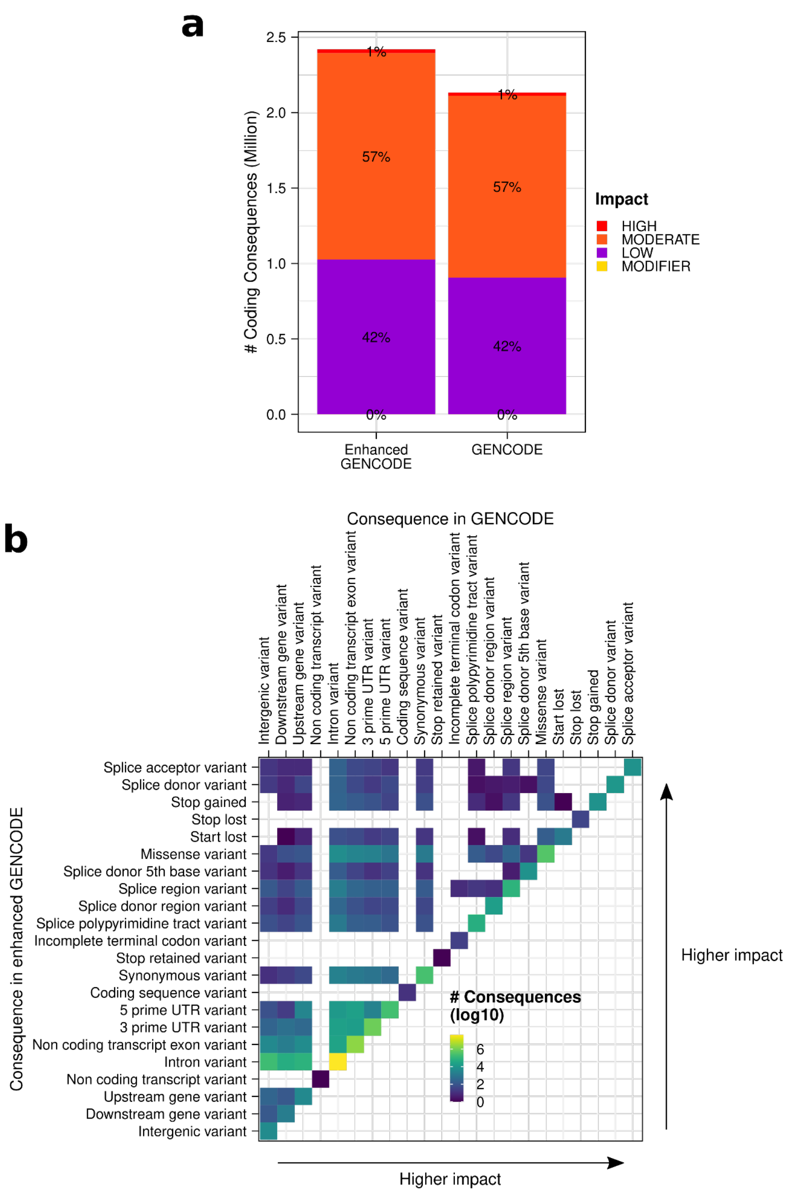


**Supplementary Fig. S13: Variant effect prediction in GENCODE and Enhanced Gencode.**
**a,** Distribution of the variant impact on all the impacted transcripts in each annotation.
**b,** For a given variant, changes in the most severe consequences predicted across all impacted transcripts between Enhanced GENCODE (left) and GENCODE (top). Consequence severity predictions are ordered according to the VEP nomenclature.

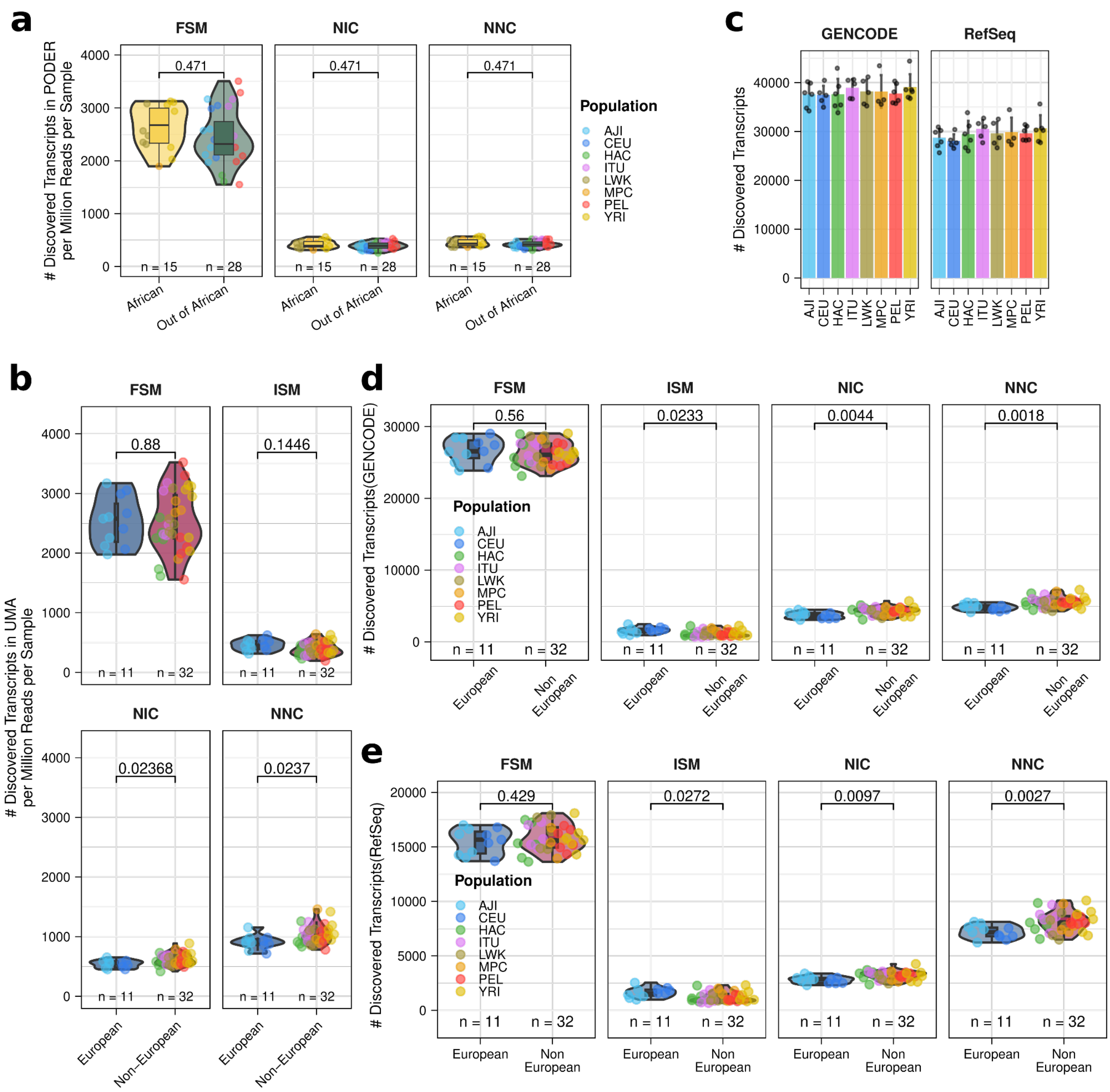


**Supplementary Fig. S14: Gene annotations are robustly European-biased.**

**a,** Number of PODER transcripts discovered per sample normalized by sequencing depth between African vs. Out of African populations by structural category (t-test, FDR adjusted by Benjamini–Hochberg).

**b,** Number of UMA transcripts (before filtering to obtain PODER; see Supplementary Fig. S4a) discovered per sample normalized by sequencing depth between European vs. non-European populations by structural category (t-test, FDR adjusted by Benjamini–Hochberg).

**c,** Number of unique transcripts discovered per sample by ESPRESSO when using (left) GENCODE and (right) RefSeq as references by downsampling to minimum read depth (8.6 million reads; ANOVA Discovered Transcripts ~ population, n.s.). Bar height represents the mean and error bars represent the standard deviation.

**d,** Number of transcripts discovered when using GENCODE as a reference per sample and downsampling to equal read depths between European vs. non-European populations by structural category (t-test, FDR adjusted by Benjamini–Hochberg).

**e,** Number transcripts discovered per sample when using RefSeq as a reference downsampling to equal read depths between European vs. non-European populations by structural category (t-test, FDR adjusted by Benjamini–Hochberg).

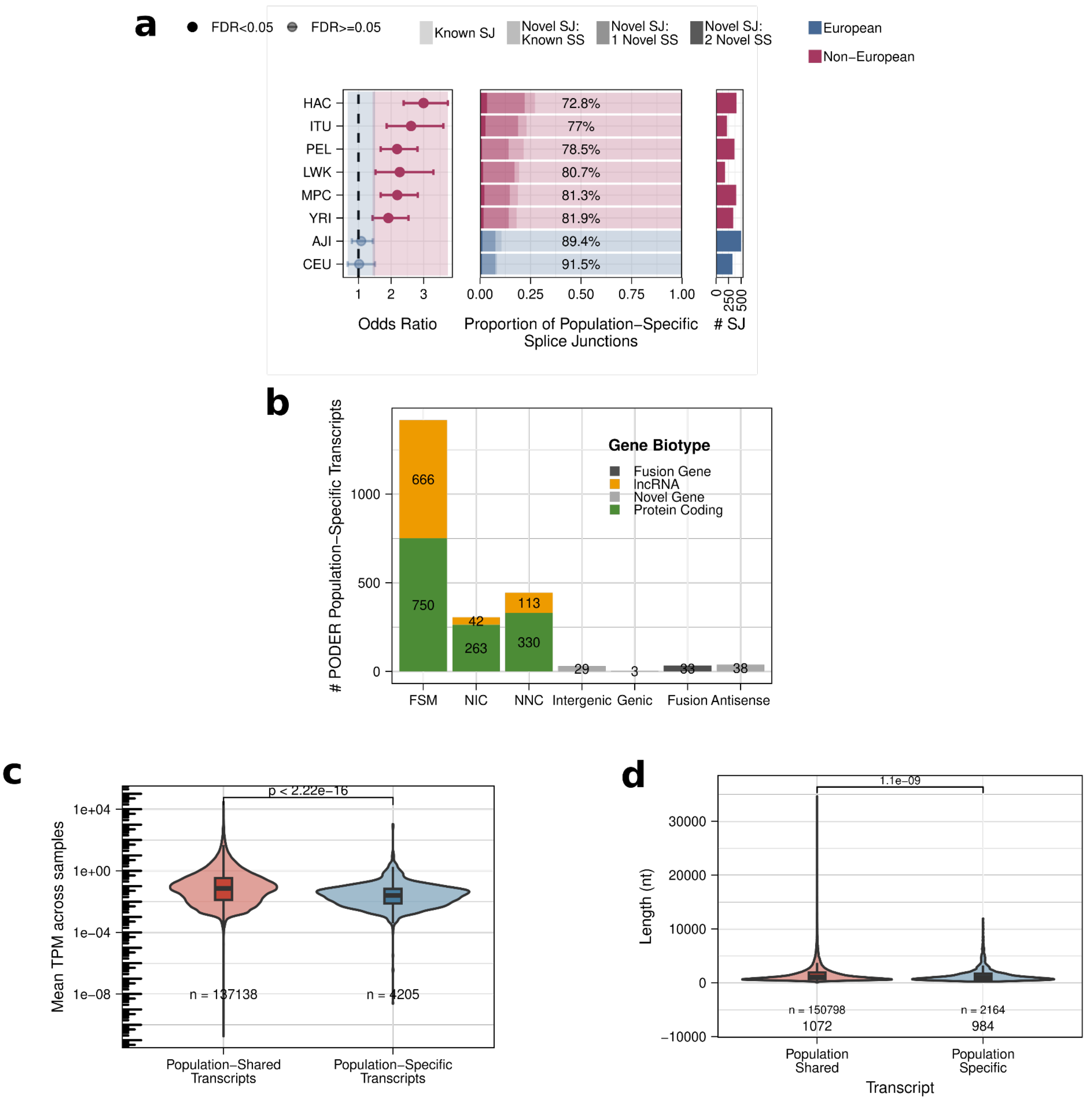


**Supplementary Fig. S15: Non-European population-specific discovered transcript and splice junction characteristics**

**a,** (Left) Enrichment in novel splice junctions of population specific splice junctions per population. (Middle) Proportion of population-specific splice junctions by annotation status. (Right) Number of population-specific splice junctions (SJ) per population.

**b,** Number of PODER population-specific transcripts by structural category and gene biotype. Novel gene category groups intergenic, antisense, and genic.

**c,** Mean transcript expression (TPM) across samples of population-shared and specific transcripts. Wilcoxon's rank-sum test.

**d,** Length in nucleotides (nt) of population-shared and specific transcripts.

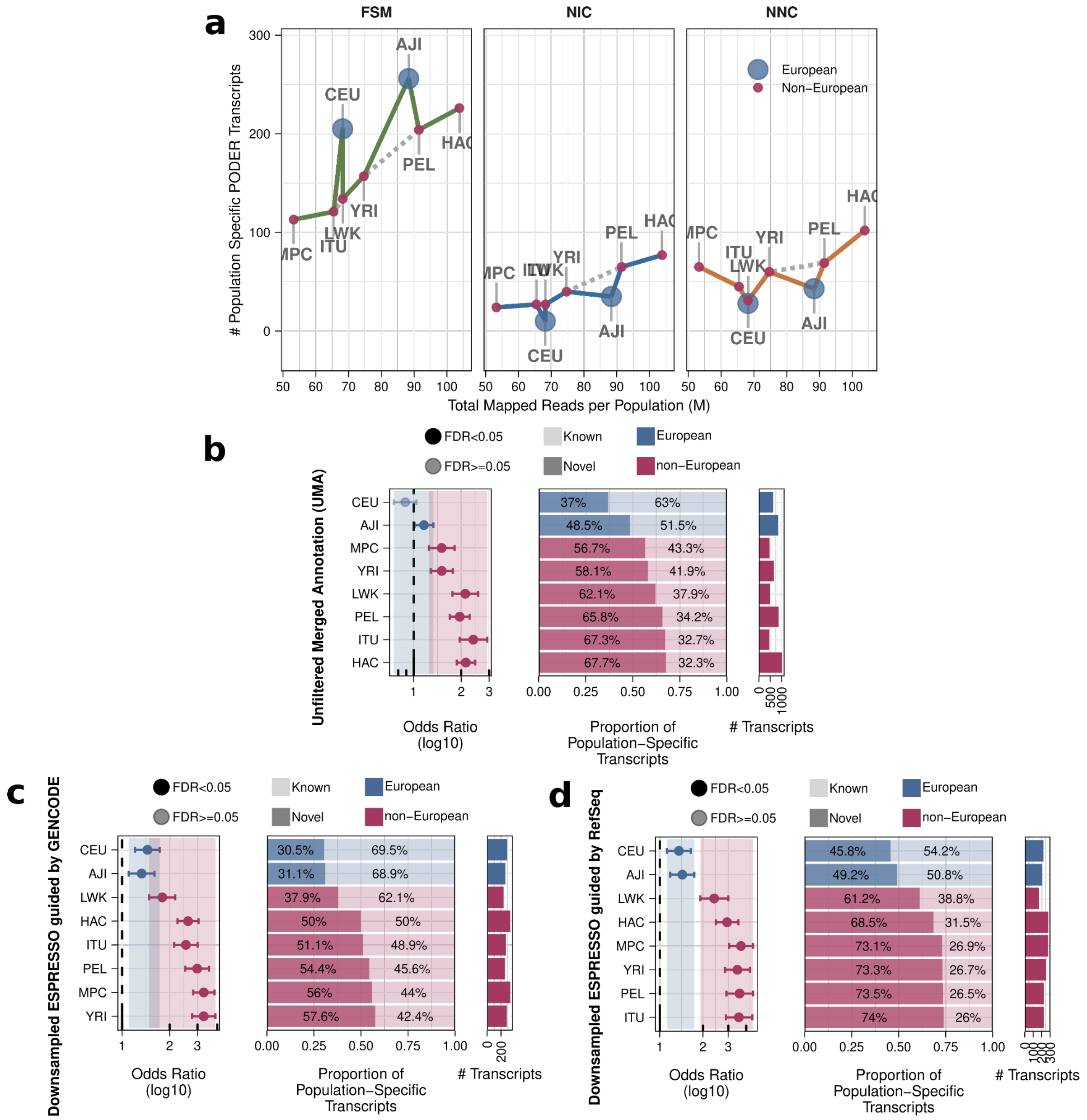


**Supplementary Fig. S16: Non-European population-specific discovered transcripts are enriched in novel**

**a,** Number of population-specific discovered transcripts from PODER by the total number of mapped reads per population. Lines colored by transcript structural category. Markers colored by whether the population is European or non-European.

**b,** (Left) Enrichment in novel transcripts of population specific transcripts from UMA (before filtering to obtain PODER; see Supplementary Fig. S4a) per population. (Middle) Proportion of population-specific transcripts by annotation status. (Right) Number of population-specific transcripts per population.

**c,** (Left) Enrichment in novel transcripts of population specific transcripts when using GENCODE as a reference downsampling to equal read depths per population. (Middle) Proportion of population-specific transcripts by annotation status. (Right) Number of population-specific transcripts per population.

**d,** (Left) Enrichment in novel transcripts of population specific transcripts when using RefSeq as a reference downsampling to equal read depths per population. (Middle) Proportion of population-specific transcripts by annotation status. (Right) Number of population-specific transcripts per population.

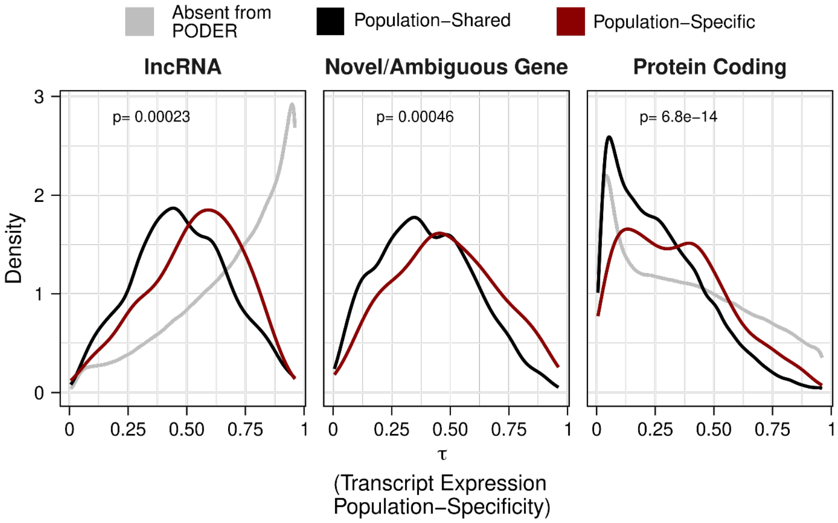


**Supplementary Fig. S17: Population-specific transcripts are supported at the expression level in the external MAGE dataset.**

Density of tau values showing transcript population-specificity in MAGE stratified by gene biotype (novel genes have an unknown biotype). Different colors show if transcripts are population-shared, population-specific, or are absent from PODER and therefore were not assessed. MAGE data quantified using Enhanced GENCODE. Wilcoxon's rank-sum test between population-shared and population-specific by gene biotype.

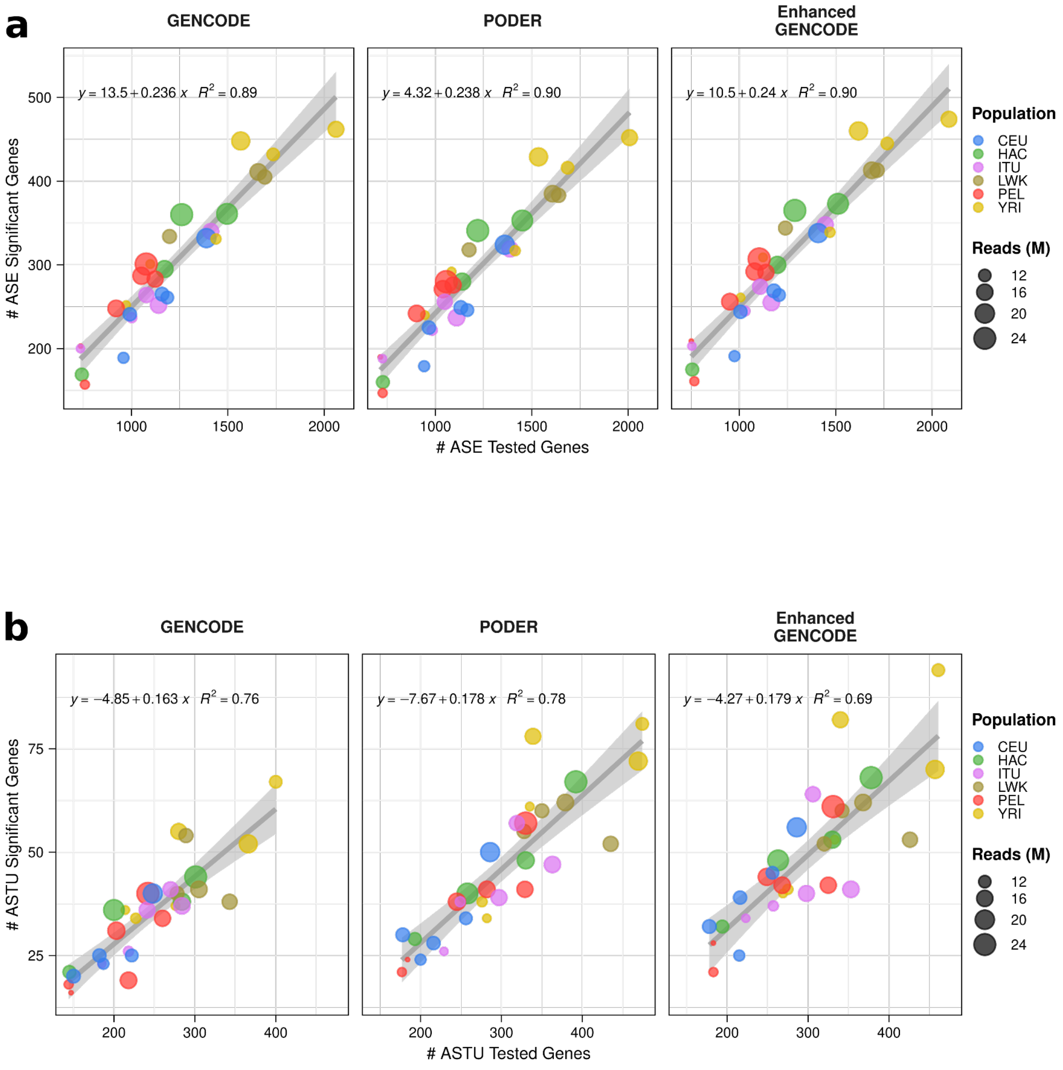


**Supplementary Figure S18: Relationship between number of testable and significant genes in allele-specific analyses.** **a-b,** Number of tested and significant (FDR < 0.05) genes for **a,** ASE; **b,** ASTU using different annotations.

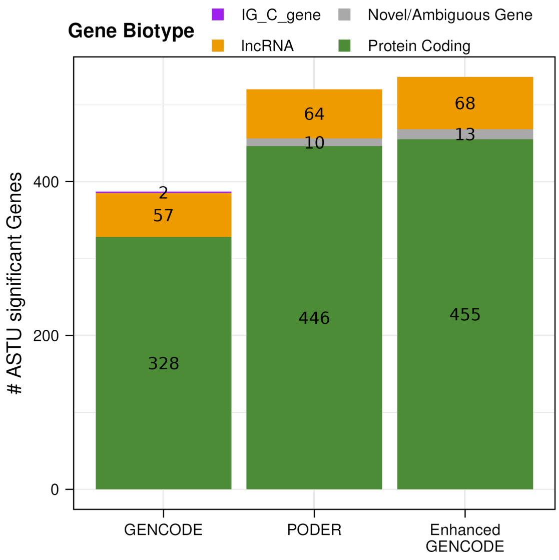


**Supplementary Figure S19: Most tested and significant ASTU genes are protein coding.** Number of ASTU genes by gene biotype using GENCODE, PODER, and Enhanced GENCODE.

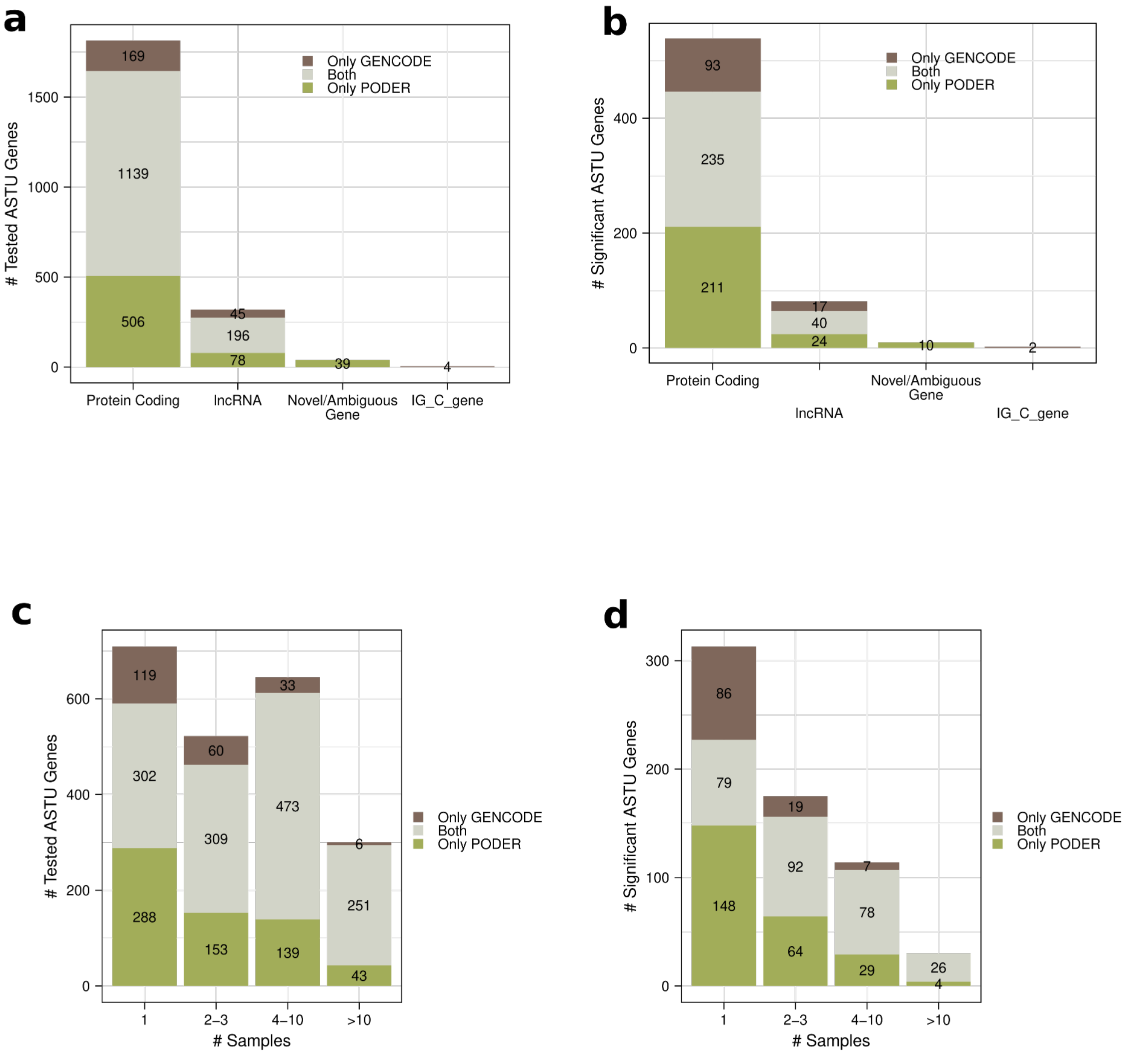


**Supplementary Figure S20: Sample and annotation sharing of ASE and ASTU tested genes.**
**a-b,** Number of **a,** tested ASTU genes **b,** significant ASTU genes by gene biotype.
**c-d,** Number of **c,** tested ASTU genes **d,** significant ASTU genes by the number of samples they were found in and how they are shared between annotations.

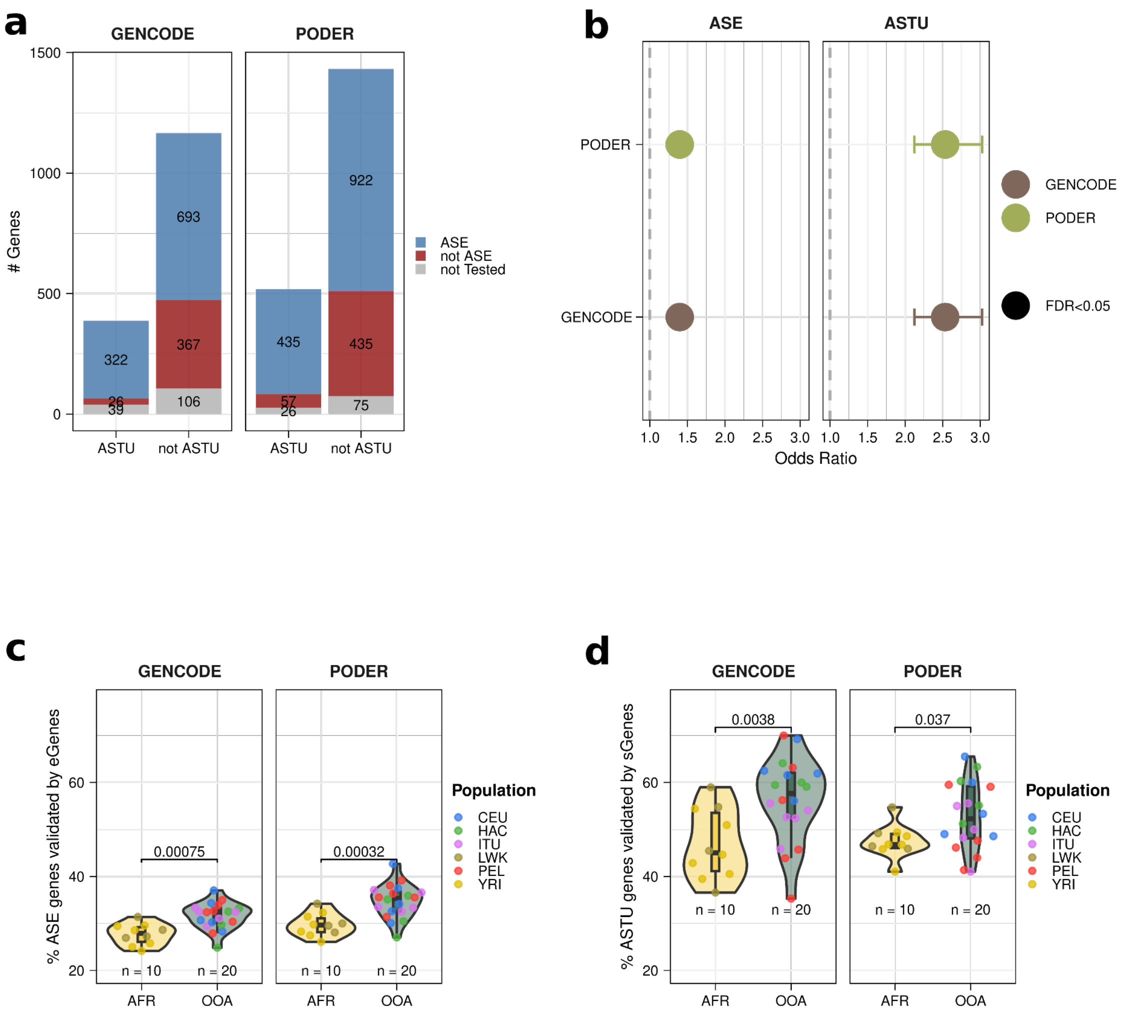


**Supplementary Figure S21: Interpretation of ASE and ASTU genes.**
**a,** Most ASTU genes are also ASE genes. Number of ASTU genes which are also ASE genes, split by GENCODE and PODER.
**b,** ASE and ASTU genes are enriched in GTEx genes with known eQTLs and sQTLS from LCLs, respectively. Odds ratio for enrichment of GTEx eGenes and sGenes for ASE and ASTU genes respectively, split by GENCODE and PODER (Fisher's exact test).
**c,** African ASE genes yield more potentially novel eGenes. Percentage of ASE genes per sample that are also eGenes; split by African and OOA (Wilcoxon rank-sum test).
**d,** African ASTU genes yield more potentially novel sGenes. Percentage of ASTU genes per sample that are also sGenes; split by African and OOA (Wilcoxon rank-sum test).

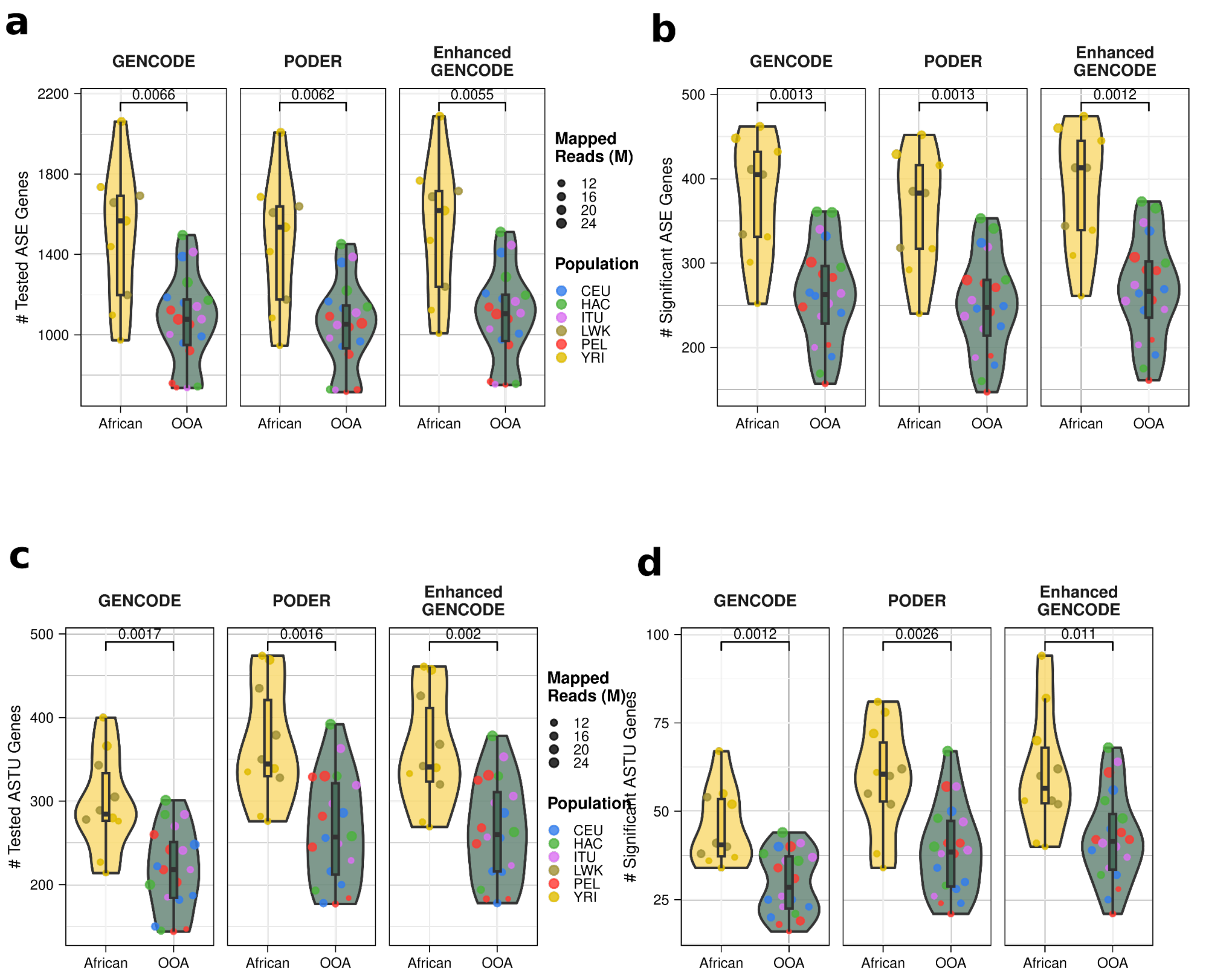


**Supplementary Figure S22: Africans have more testable and significant ASE and ASTU genes.**
**a-d,** Number of **a,** tested ASE genes **b,** significant ASE genes **c,** tested ASTU genes **d,** significant ASTU genes per sample, split by African and OOA, using GENCODE, PODER, and Enhanced GENCODE (t-test).

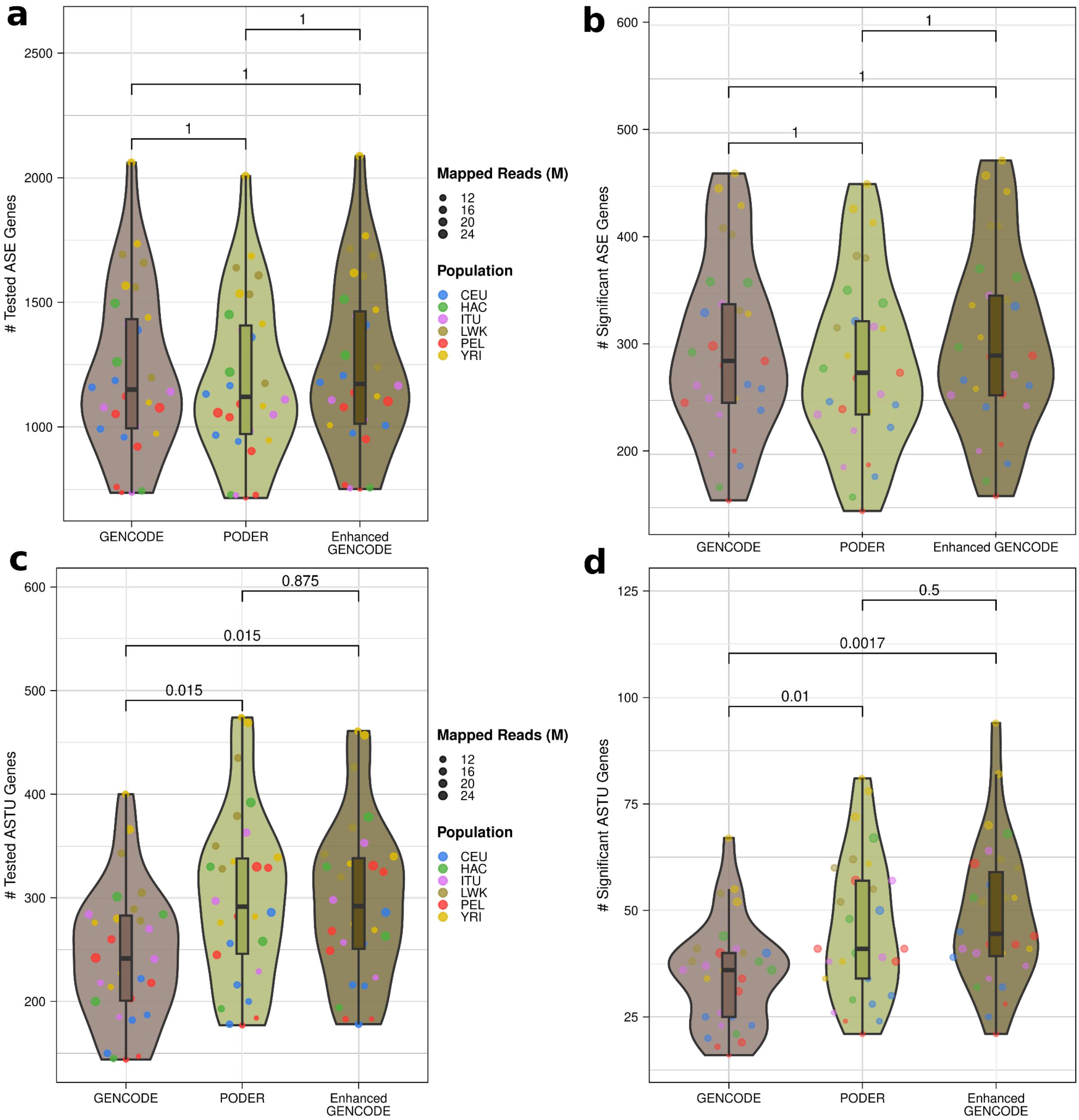


**Supplementary Figure S23: Novel PODER transcripts increases discovery of ASTU genes**.
**a-d,** Number of **a,** tested ASE genes **b,** significant ASE genes **c,** tested ASTU genes **d,** significant ASTU genes per sample, using GENCODE, PODER, and Enhanced GENCODE (t-test).

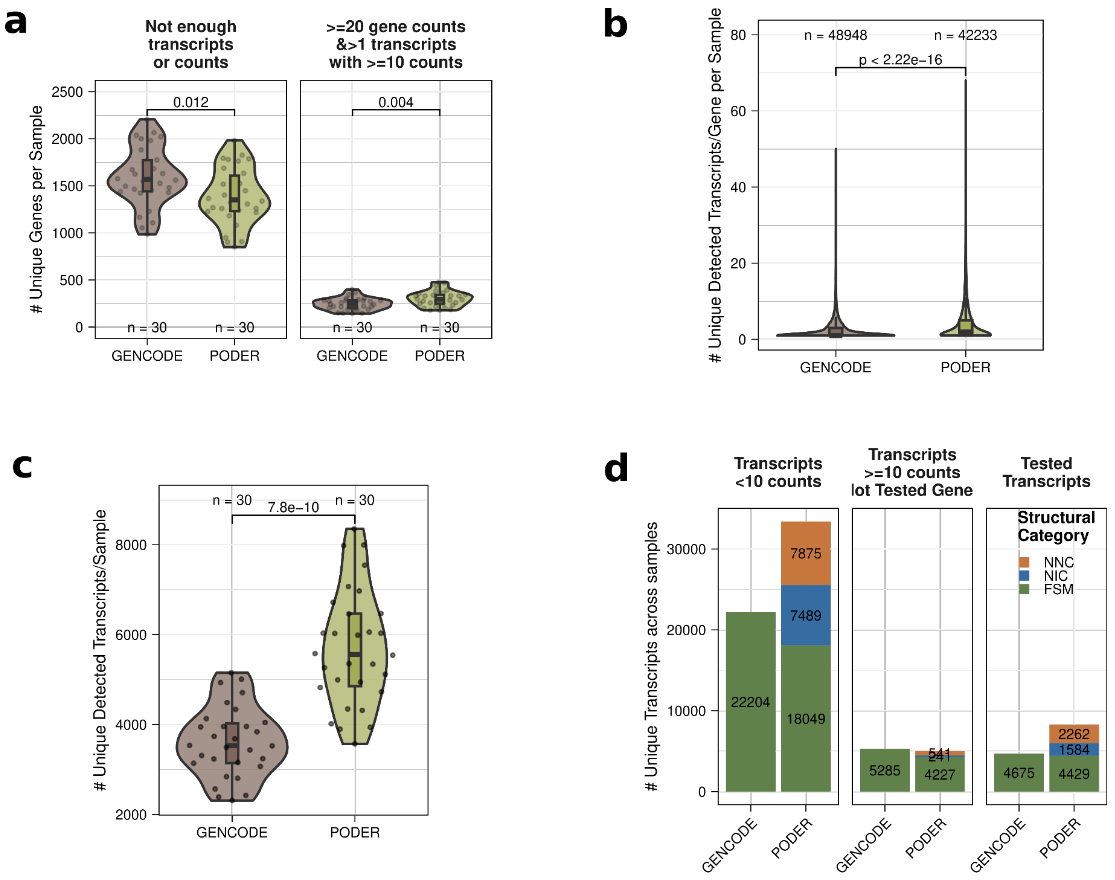


**Supplementary Figure S24: The addition of novel transcripts to the annotation increases testable ASTU genes.**
**a,** Number of unique genes per sample not testable (left) versus number of testable genes per sample (right) using GENCODE or PODER.
**b,** Overall number of detected transcripts per gene in each sample using GENCODE or PODER.
**c,** Overall number of detected transcripts in each sample using GENCODE or PODER.
**d,** Number and novelty of transcripts passing or failing various filters required for ASTU testing.

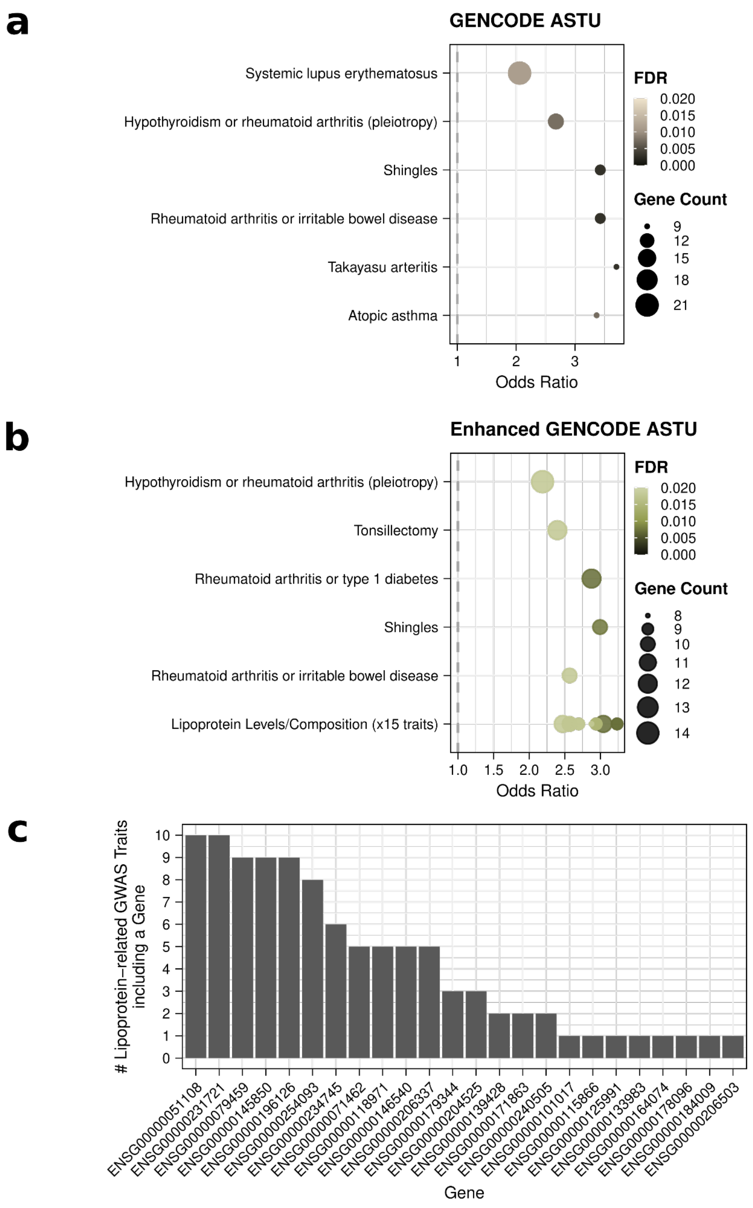


**Supplementary Figure S25: ASTU genes are enriched for GWAS genes harboring or close to variants from ancestry-divergent traits.**

**a,** GENCODE ASTU significant genes enrichment in GWAS hit associated genes (cutoff FDR=0.02).

**b,** Enhanced GENCODE ASTU significant genes enrichment in GWAS hit associated genes (cutoff FDR=0.02).

**c,** Number of lipoprotein-related GWAS traits hits in PODER containing a gene that is ASTU significant.

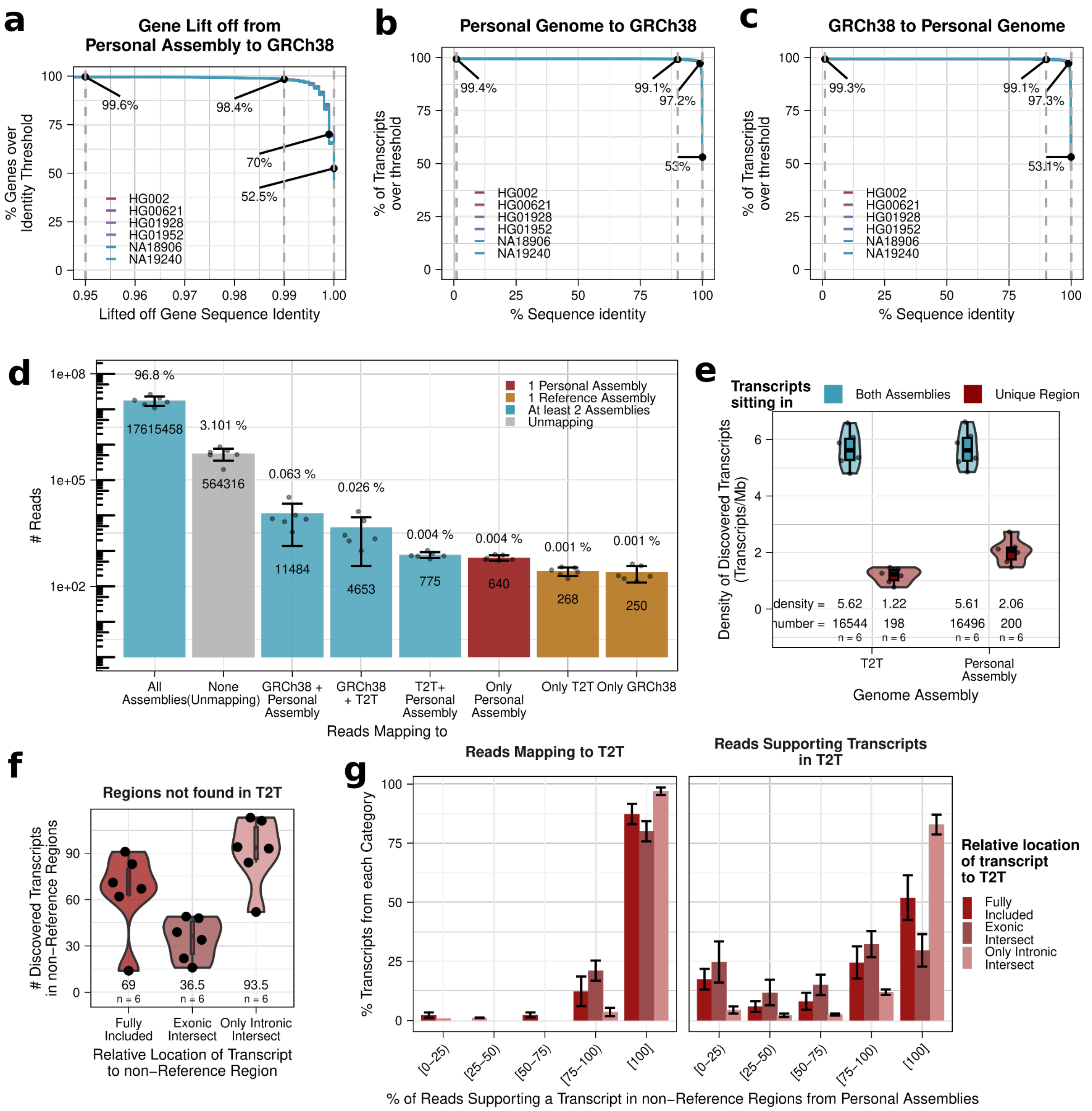


**Supplementary Figure S26: Personal genome regions not found in GRCh8 or T2T harbour some lncRNA or protein-coding genes.**

**a,** Percentage of lifted off genes from personal assemblies to GRCh38 over a certain sequence identity threshold with GRCh38. Numbers indicate (left to right) % genes with at least 95%, 99%, 99.9% and 100% sequence identity. **b-c,** Percentage of personal assembly (or GRCh38) discovered transcripts with a BLAST match against transcripts discovered in GRCh38 (or personal assembly) over the threshold (x-axis) of sequence identity percentage with their best match (i.e. 99.1% of transcripts discovered in the personal assemblies have at least a 90% sequence identity with a transcript discovered in GRCh38). Different colors represent transcripts from different personal assemblies. **d,** Mean number of reads by the combinations of assemblies where they map to. **e,** Density of transcripts in transcripts/Mb of DNA sequence discovered in each genome assembly by if transcript is located in a region shared across genome assemblies or intersecting an assembly-specific region. **f,** Number of discovered transcripts intersecting non-reference regions by their relative position to non-reference regions. Number shows the median. **g,** Most reads that discover a transcript in personal assemblies also (left) map to and (right) are used to discover transcripts in T2T. Mean percentage of transcripts from non-reference regions, binned by their percentage of supporting reads (on the left, for mapping; on the right, for supporting the discovery of a transcript in T2T). Error bars show standard deviations.

**a**

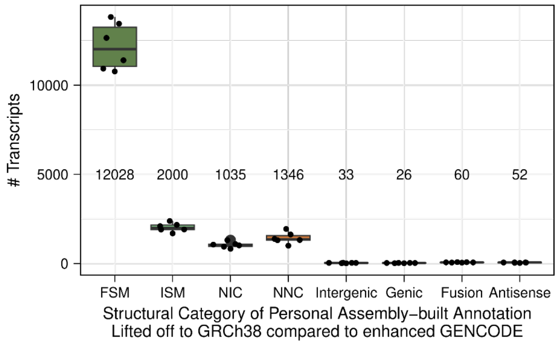


**b**

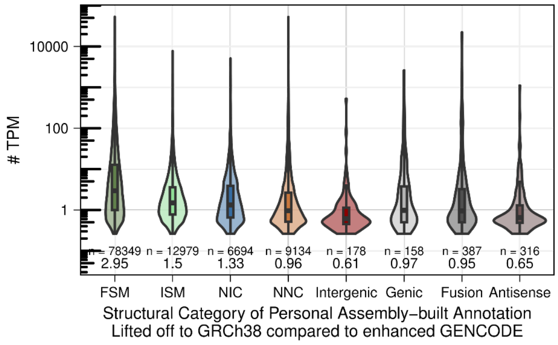


**Supplementary Figure S27: Comparison of transcripts discovered in personal assemblies with respect to Enhanced GENCODE.**

**a,** Number of transcripts discovered in each personal assembly by structural category after lifting over GRCh38 with respect to Enhanced GENCODE. **b,** Transcript expression in TPM of lifted over transcripts by structural category. Bottom numbers are medians.

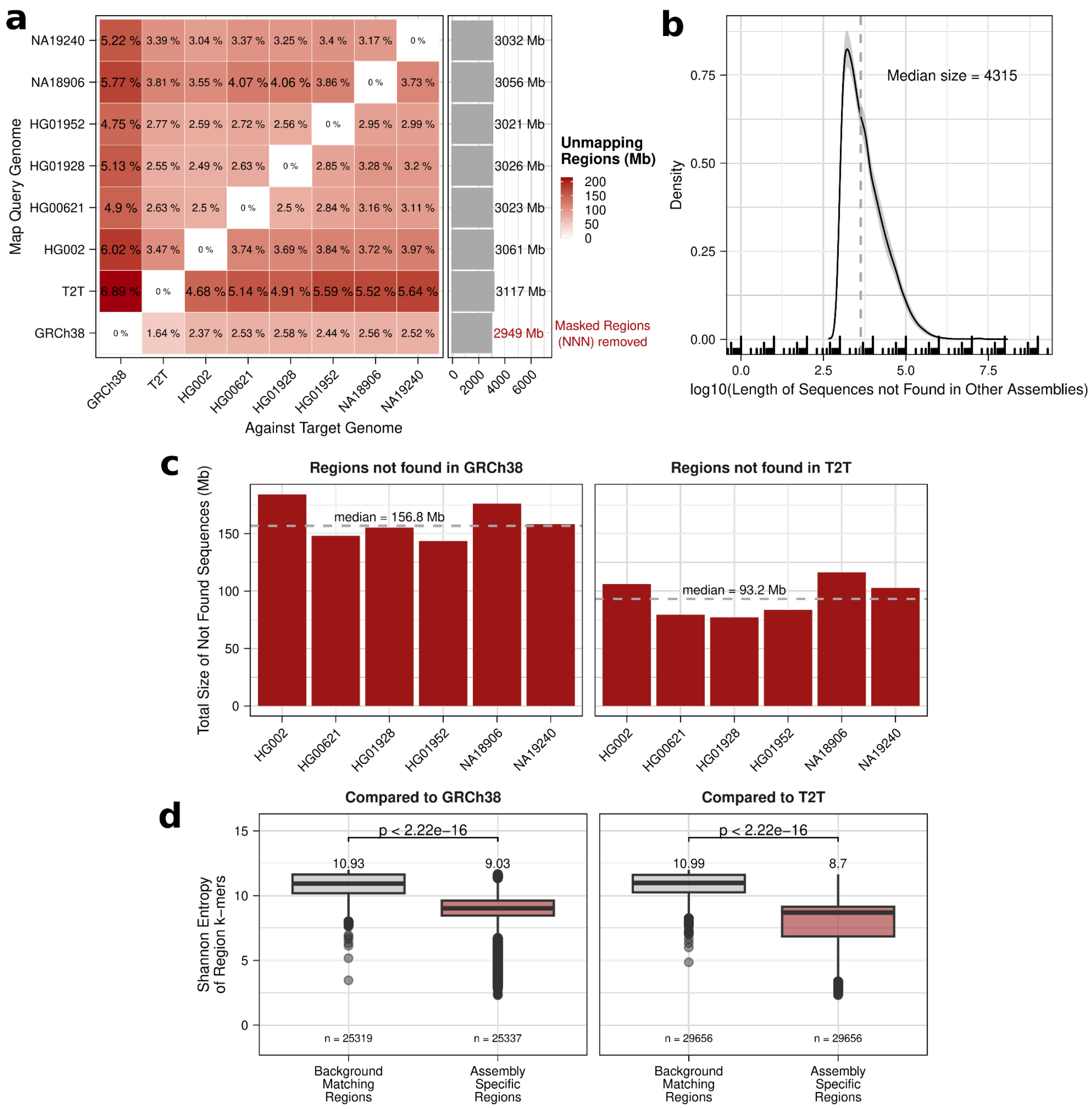


**Supplementary Figure S28: Non-reference regions from personal assemblies that are short and with low complexity.** **a,** Percentage of query genome not found (unmapping) in target genome (left), referred to as non-reference regions, and number of Mb in query genome. Percentages given with respect to target genome length. **b,** Mean size of non-reference regions across personal assemblies with respect to GRCh38, shade represents standard deviation. **c,** Total length of non-reference regions per personal assembly with respect to GRCh38 and T2T. **d,** Shannon entropy of non-reference regions k-mers and length-matched background regions from the rest of the assembly. Wilcoxon's rank-sum test.
